# Structural and biochemical analyses of selectivity determinants in chimeric *Streptococcus* Class A sortase enzymes

**DOI:** 10.1101/2021.09.19.461001

**Authors:** Melody Gao, D. Alex Johnson, Isabel M. Piper, Hanna M. Kodama, Justin E. Svendsen, Elise Tahti, Brandon Vogel, John M. Antos, Jeanine F. Amacher

**Author notes:** Corresponding Author: Jeanine Amacher, Department of Chemistry, Western Washington University, 516 High St – MS9150, Bellingham, WA, 98225. These authors contributed equally to this work (Melody Gao, D. Alex Johnson).

## Abstract

Sequence variation in related proteins is an important characteristic that modulates activity and selectivity. An example of a protein family with a large degree of sequence variation is that of bacterial sortases, which are cysteine transpeptidases on the surface of gram-positive bacteria. Class A sortases are responsible for attachment of diverse proteins to the cell wall to facilitate environmental adaption and interaction. These enzymes are also used in protein engineering applications for sortase-mediated ligations (SML) or *sortagging* of protein targets. We previously investigated SrtA from *Streptococcus pneumoniae*, identifying a number of putative β7-β8 loop-mediated interactions that affected *in vitro* enzyme function. We identified residues that contributed to the ability of *S. pneumoniae* SrtA to recognize several amino acids at the P1’ position of the substrate motif, underlined in LPXT<u>G</u>, in contrast to the strict P1’ Gly recognition of SrtA from *Staphylococcus aureus*. However, motivated by the lack of a structural model for the active, monomeric form of *S. pneumoniae* SrtA, here, we expanded our studies to other *Streptococcus* SrtA proteins. We solved the first monomeric structure of *S. agalactiae* SrtA which includes the C-terminus, and three others of β7-β8 loop chimeras from *S. pyogenes* and *S. agalactiae* SrtA. These structures and accompanying biochemical data support our previously identified β7-β8 loop-mediated interactions and provide additional insight into their role in Class A sortase substrate selectivity. We argue that a greater understanding of individual SrtA sequence and structural determinants of target selectivity can facilitate the design or discovery of improved sortagging tools.

## Introduction

Class A sortases are enzymes located on the surface of gram-positive bacteria that attach proteins to the cell wall. Sortase-mediated protein display allows bacteria to interact with their environments, e.g., with proteins for bacterial adhesion and/or acquisition of nutrients, and can include pathogenic factors that enable the bacteria to infect host organisms.^1,2^ The catalytic mechanism of sortases involves the recognition and cleavage of a specific sequence, followed by ligation of an incoming amine nucleophile.^1–3^ This reactivity has also been harnessed for protein engineering applications, and sortases have emerged as powerful tools for the post-translational derivatization of protein targets with various non-native modifications.^3^ The traditional recognition motif of Class A sortase (or SrtA) proteins, which is found within the cell wall sorting signal (CWSS) of gram positive bacteria, is the sequence LPXTG (where X = any amino acid, and L=P4, P=P3, X=P2, T=P1, and G=P1’). This sequence is recognized by all Class A sortases investigated to date, however, other recognition sequences have been identified or engineered for several SrtA proteins in the last decade, greatly increasing the potential for sortase-mediated ligation (SML), or *sortagging*, applications.^4–12^

Despite a relatively large degree of sequence variation amongst the hundreds of identified SrtA proteins in bacteria, these cysteine transpeptidases contain a conserved catalytic triad, consisting of His, Cys, and Arg residues.^1,2,13^ The most well studied Class A sortase is that from *Staphylococcus aureus* (saSrtA), which continues to see frequent use in sortagging applications.^3^ As of 2019, there were approximately 10 known structures of Class A sortases, with several being of saSrtA.^13^ Overall, the sortase fold, consisting of a closed 8-stranded β-barrel architecture, is conserved in all structures of SrtA proteins solved to date; however, there are variations consistent with the degree of sequence differences.^2^ For example, between saSrtA and *Streptococcus pyogenes* SrtA (spySrtA), there are a number of unique structural characteristics that affect enzyme function (**Figure 1**). Specifically, saSrtA requires a Ca^2+^ cofactor and its β7-β8 loop near the active site contains an additional 5 residues and a Trp (W194) which dramatically affects activity (**Figure 1a**).^12,14^ All structural comparisons with saSrtA will use the peptidomimetic-bound structure (PDB ID 2KID) as this is the only one to our knowledge of saSrtA in the active state.^15–19^ Previous work shows that allosteric activation, driven by Ca^2+^ binding, affects several structural features near the active site, including the relative conformation and/or location of the β6-β7 and β7-β8 loops.^15–20^ In the case of spySrtA, a partially helical C-terminal extension of 24 residues is evident in the reported crystal structure that is absent in SaSrtA (**Figure 1**). A detailed description of how each of these features determine target recognition and selectivity remains incomplete, particularly for the *Streptococcus* SrtA proteins. Unique features of *B. anthracis* SrtA (baSrtA) have also been previously described, e.g., regulation of enzymatic activity by an N-terminal appendage as well as a disordered-to-ordered transition in the β7-β8 loop upon ligand binding.^21^

**Figure 1.**
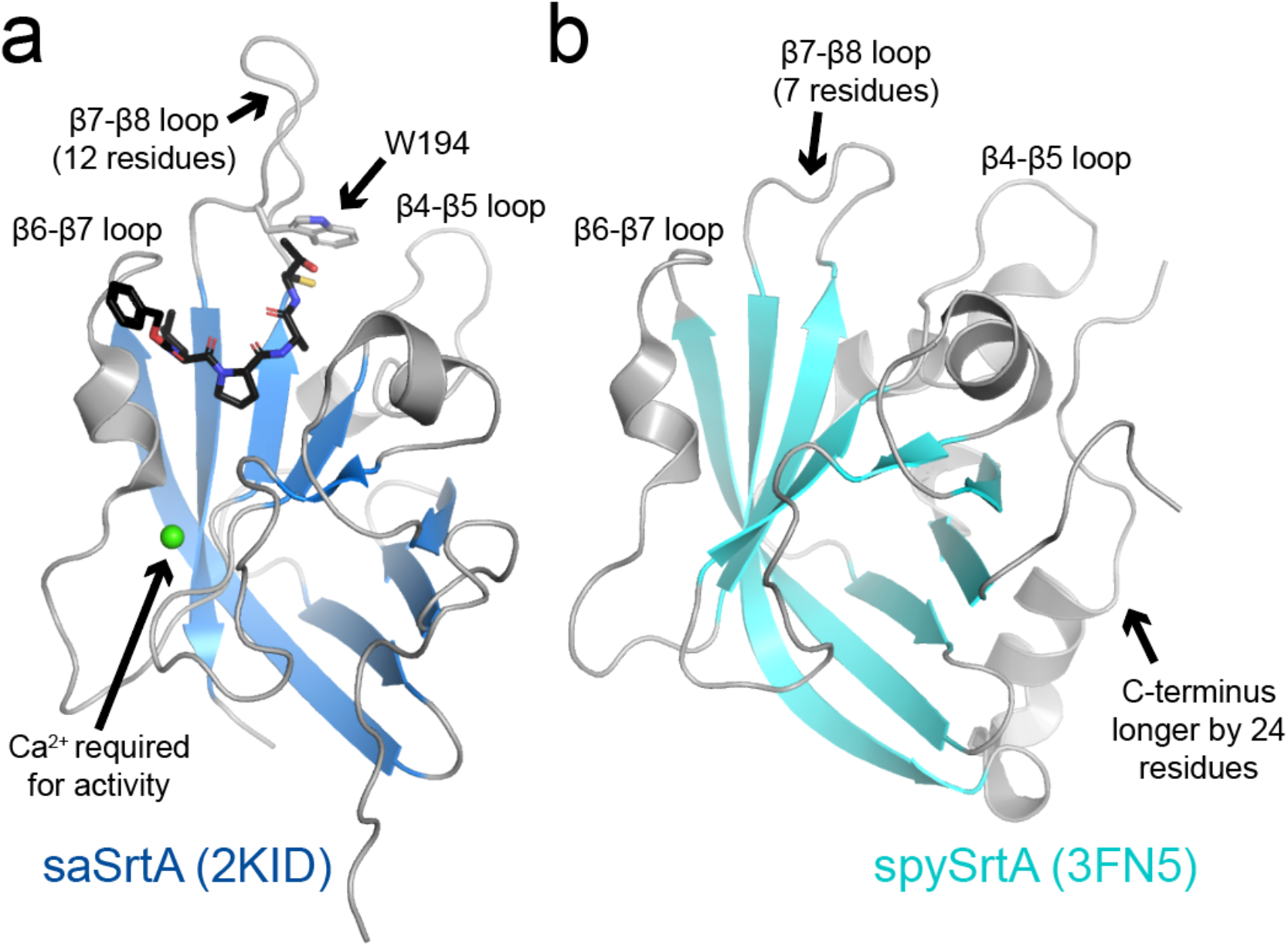
Differences between *Staphylococcus* and *Streptococcus* SrtA proteins. The SrtA proteins are in cartoon representation, with the conserved 8-stranded antiparallel β-sheet that defines the sortase fold colored as labeled. *S. aureus* SrtA (saSrtA, PDB ID 2KID) is bound to the peptidomimetic, LPAT*, in black sticks and colored by heteroatom (O=red, N=blue, S=yellow), (**a**). The arrows indicate differences with *S. pyogenes* SrtA (spySrtA, 3FN5) (**b**). Specifically, Ca^2+^ is required for saSrtA activity, the β7-β8 loop contains an additional 5 residues and contains a Trp (W194) that dramatically affects activity, and the spySrtA protein is 24 residues longer than saSrtA.

A number of protein families use specificity-determining loops to encode differing target selectivity amongst members. Classic examples include kinases and serine proteases.^22–28^ Specific regions of the activation loop of kinases contribute to substrate specificity by directly interacting with amino acids adjacent to the phosphorylation site.^25,28^ In serine proteases, substitution of two conserved surface loops (9 residues total) efficiently converts selectivity of trypsin to that of the related enzyme chymotrypsin.^22,23^ There are also examples in scaffolding domains, including SH2 and SH3 domains, where conserved loops interact directly with the peptide and determine the selectivity of both SH2 (the EF and BG loops) and SH3 (the RT and n-Src loops) domains.^29–37^ Work from ourselves and others strongly indicates that Class A sortases are another protein family that exhibits functionally relevant sequence variation in specificity-determining loops.^12,38,39^

In our previous work, we investigated the selectivity determinants of *Streptococcus pneumoniae* SrtA (spSrtA) at the P1’ position of the CWSS.^12^ We found that the sequence of the β7-β8 loop dramatically affects enzyme activity and selectivity at this substrate position.^12^ Because spSrtA crystallizes as a domain-swapped dimer, which is enzymatically inactive in our hands, we used previously published Class A sortase structures to investigate the stereochemistry of our biochemical results.^12,40,41^ Now, we investigate two additional sortases, those from *S. pyogenes* (spySrtA) and *S. agalactiae* (sagSrtA), to see if the β7-β8 loop has broad effects on enzyme function and target recognition for *Streptococcus* Class A sortases.

We find that the β7-β8 loop affects spySrtA and sagSrtA in a manner consistent with that of *S. pneumoniae* SrtA. To investigate how the β7-β8 loop sequence affects each protein, we created a series of chimeric enzymes, swapping the loop sequences from several of those previously studied.^11,12^ As seen previously, while some loop sequences hinder enzyme activity in our FRET-based assay, others improve target substrate cleavage, which is the presumed rate-limiting step of the sortase-catalyzed transpeptidation reaction.^42^ Here, we also use X-ray crystallography to look at the stereochemistry of both spySrtA and sagSrtA β7-β8 chimeric proteins. Finally, we use mutagenesis, structural, and sequence analyses to investigate conserved characteristics in the β7-β8 loops from *Streptococcus* SrtA proteins. Taken together, these analyses provide new insights on the role of conserved loops near the active site of *Streptococcus* Class A sortases.

## Results

### Enzyme assays of wild-type S. pyogenes and S. agalactiae SrtA proteins

Based on our results using spSrtA, we designed a number of β7-β8 loop chimeras using spySrtA and sagSrtA as the “scaffolds.” The wild-type sequences used were spySrtA_82-249_ (PDB ID 3FN5), sagSrtA_79-238_ (the sequence crystallized previously, in PDB ID 3RCC), or sagSrtA_79-247_, which includes the final nine C-terminal residues of sag SrtA based on UniProt ID SRTA_STRA3.^43–45^ For simplicity, we will refer to these as: spySrtA, sagSrtA_238_, and sagSrtA_247_. The β7-β8 loop sequences of these wild-type constructs were as follows: spySrtA (sequence: <u>C</u>TDIEATE<u>R</u>, the catalytic Cys and Arg are included and underlined as reference points for the loop boundaries) and sagSrtA (<u>C</u>TDPEATE<u>R</u>). Notably, the β7-β8 loops of spySrtA and sagSrtA differ at only one position, 3 residues C-terminal to the catalytic Cys, which we will refer to as β7-β8^+3^. The β7-β8^+3^ residue is Ile in spySrtA and Pro in sagSrtA. The wild-type sequences of spySrtA and sagSrtA_247_ are overall 65% identical (**Figure 2a**), which is consistent with relative sequence identities amongst other representative *Streptococcus* Class A sortases (**Figure S1**).

**Figure 2.**
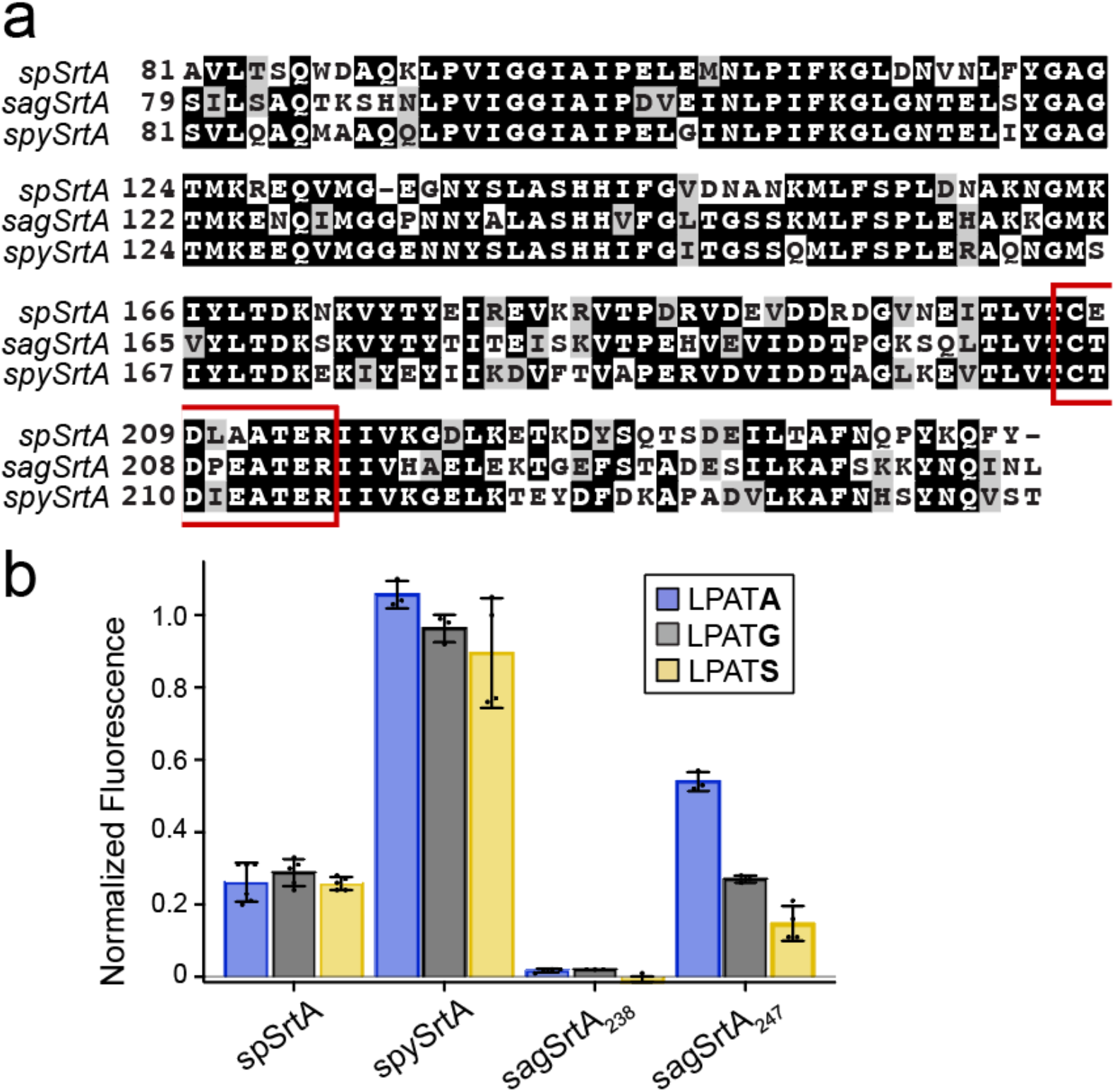
Biochemical characteristics of SrtA enzymes from *S. agalactiae* and *S. pyogenes*. (**a**) Sequence alignment of the extracellular regions of the *S. pneumoniae* SrtA (spSrtA), *S. agalactiae* SrtA (sagSrtA) and *S. pyogenes* SrtA (spySrtA) proteins. Sequences were aligned using T-coffee and visualized with Boxshade. The β7-β8 loop residues are indicated with a red box. (**b**) Comparison of substrate selectivity for wild-type spSrtA, spySrtA, sagSrtA_238_ (the crystallized construct reported in PDB ID 3RCC), and spySrtA_247_. Substrate cleavage was monitored via an increase in fluorescence at 420 nm from reactions of the fluorophore-quencher probes Abz- LPATGG-K(Dnp), Abz-LPATAG-K(Dnp), and Abz-LPATSG-K(Dnp) (represented as LPAT**G**, LPAT**A**, and LPAT**S**) in the presence of excess hydroxylamine. Bar graphs represent mean normalized fluorescence (± standard deviation) from at least three independent experiments at the 2 h reaction timepoint, as compared to saSrtA and the peptide LPAT**G**. The spSrtA data was previously published and is shown for comparison.^12^ Averaged assay values and standard deviations for spySrtA, sagSrtA_238_, and sagSrtA_247_ are in **Table S1**.

In order to assess relative activity and selectivity, we used a FRET-based enzyme assay involving synthetic peptide substrates. This assay utilizes well-established FRET quencher probes consisting of a substrate sequence with an N-terminal 2-aminobenzoyl fluorophore (Abz) and C-terminal 2,4-dinitrophenyl (Dnp) quencher.^12,14,46,47^ For all assays, fluorescence was monitored for 2 h at room temperature and analyzed relative to a benchmark reaction consisting of wild-type saSrtA and the Abz-LPATGG-K(Dnp) peptide.^12^ For simplicity, we will remove the “Abz-” and “G-K(Dnp)” from peptide names hereafter, as they are not a part of the CWSS (e.g., Abz- LPATGG-K(Dnp) will be referred to as LPAT**G**). Additional experimental details are provided in the Materials and Methods and all averaged assay data and standard deviation values are in **Table S1**. All sortase enzymes were expressed and purified as previously described and as in the Materials and Methods.^12^ Purity was assessed by SDS-PAGE and monomeric protein fractions were pooled following size exclusion chromatography, as previously described.^12^

With the necessary materials in hand, we first evaluated the reactivity of wild-type spySrtA, sagSrtA_238_, and sagSrtA_247_ proteins with peptides differing only at the P1’ position (indicated in **bold**): LPAT**A**, LPAT**G**, and LPAT**S** (**Figure 2b**). As our data shows, spySrtA was quite active, and exhibited robust reactivity that was comparable to the benchmark saSrtA/LPAT**G** reaction. This protein also exhibited comparable reactivity at the 2 h reaction endpoint with G-, S-, and A-containing peptides (**Figure 2b**). Activity for spySrtA was also markedly higher than spSrtA, which is consistent with a loop interaction described in our previous work.^12^ Specifically, the β6^-2^ position in spSrtA is R184, which was found to have a negative impact on reactivity that was attributed to a putative interaction with the β7-β8^-1^ Glu of this enzyme.^12^ In spySrtA, the corresponding β6^-2^ position is T185, which likely minimizes this interaction and increases reactivity. Indeed, the spySrtA structure does not show evidence for this type of interaction (**Figure S2a**). Turning to the *S. agalactiae* constructs, sagSrtA_238_ was catalytically inactive for all peptides tested, while sagSrtA_247_ reacted with all three, albeit at a lower level than spySrtA (**Figure 2b**). SagSrtA_247_ also exhibited a preference for LPAT**A**, and we observed 50% and 72% reductions in the relative activities for the G- and S-containing peptides as compared to LPAT**A**, respectively. This is in contrast to spSrtA, which displayed almost identical relative fluorescence values after 2 h for G-, S-, and A-containing peptides (0.29 ± 0.04, 0.26 ± 0.02, and 0.26 ± 0.01, respectively) (**Figure 2b**).^12^

### The β7-β8 loop of Streptococcus SrtA proteins broadly affects enzyme activity and selectivity

Based on our previous results with spSrtA, we next wanted to substitute β7-β8 loop sequences from other SrtA proteins into spySrtA and sagSrtA_247_.^12^ We chose to substitute the SrtA β7-β8 loop sequences from: *S. aureus* (<u>C</u>DDYNEKTGVWEK<u>R</u>), *E. faecalis* (<u>C</u>GDLQATT<u>R</u>), *L. monocytogenes* (<u>C</u>DKPTETTK<u>R</u>), and *S. pneumoniae* (<u>C</u>EDLAATE<u>R</u>). These sequences were chosen due to their variable effects on spSrtA.^12^ For example, spSrtA_aureus_ (subscript denotes origin of the β7-β8 loop sequence) was relatively active and selective for a P1’ Gly residue, spSrtA_faecalis_ was relatively active and non-selective at P1’, and spSrtA_monocytogenes_ was inactive.^12^

In total we tested eight additional variants: spySrtA_aureus_, spySrtA_faecalis_, spySrtA_monocytogenes_, spySrtA_pneumoniae_, sagSrtA_aureus_, sagSrtA_faecalis_, sagSrtA_monocytogenes_, and sagSrtA_pneumoniae_. All proteins were expressed, purified and characterized as described previously and in the Materials and Methods.^12^ In general, we saw similar trends to those seen with spSrtA (**Figure 3**).^12^ For example, in the case of the *S. aureus* loop swaps we observed that both spySrtA_aureus_ and sagSrtA_aureus_ were selective for LPAT**G**, as seen with spSrtA (**Figure 3**). All constructs containing the *L. monocytogenes* also showed a preference for LPAT**G**, along with a clear reduction (2-3 fold) in activity as compared to the wild-type enzyme.

**Figure 3.**
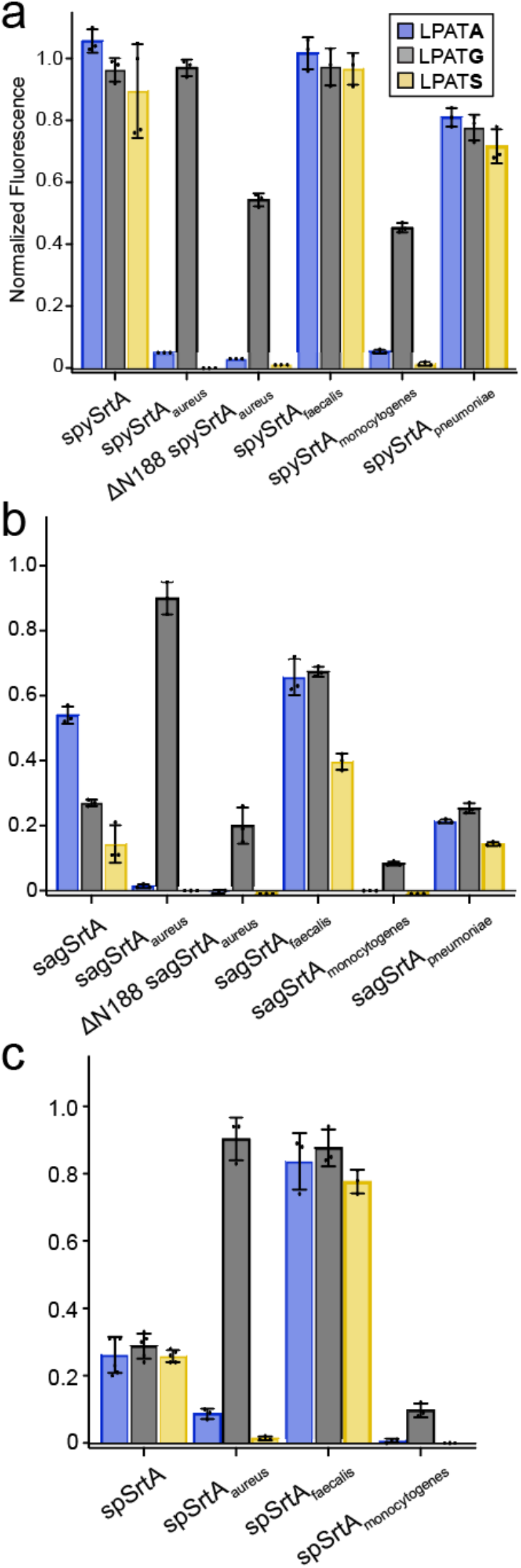
Enzyme assays of β7-β8 loop chimeras of spySrtA, sagSrtA, and spSrtA. Comparison of substrate selectivity for β7-β8 loop chimeras of (**a**) spySrtA, (**b**) sagSrtA, and (**c**) spSrtA proteins. Assays were run and data was collected as in **Figure 2**. Averaged assay values and standard deviations for spySrtA and sagSrtA variants are in **Table S1**. The spSrtA data was previously published and is shown for comparison.^12^

For the *E. faecalis* loop swaps, both the sagSrtA_faecalis_ and spySrtA_faecalis_ variants exhibited good reactivity that was generally higher than the corresponding wild-type enzyme. In contrast to spSrtA, however, the observed reactivity changes were not uniform across G-, S-, and A-containing peptides. Specifically, for sagSrtA_faecalis_ the relative activities for the LPAT**G** and LPAT**S** peptides were increased ∼2.5-fold and 2.6-fold, respectively, as compared to wild-type, whereas for LPAT**A**, it was only increased 1.2-fold (**Figure 3b**). In the case of spySrtA_faecalis_, analysis at the 2 h reaction timepoint initially suggested activity comparable to the wild-type enzyme and no preference for G-, S-, and A-containing peptides (**Figure 3a**). However, differences between spySrtA and spySrtA_faecalis_ were evident at earlier reaction time points (**Figure S3**). In particular, at early stages in the reaction (e.g. 10 min) the *E. faecalis* β7-β8 loop in spySrtA_faecalis_ appeared to significantly increase reactivity with the G- and S-containing peptides (>2-fold relative to spySrtA), and may have also had an effect on LPAT**A**, but it is inconclusive due to large error bars for early time points in this reaction (**Figure S3d**).

Finally, installation of the *S. pneumoniae* loop resulted in decreases in activity for most enzyme-substrate combinations. For spySrtA_pneumoniae_, activity was ∼20% lower than spySrtA for all peptides tested. In sagSrtA_pneumoniae_, the activity of the protein for the G- and S-containing peptides was similar to sagSrtA_247_, but was reduced ∼2.6-fold for LPAT**A** (**Figure 3b**). Focusing on spSrtA and spySrtA, comparison of the β7-β8 loop sequences revealed that while the final three positions of the 7-residue loop are identical (ATE), there are three differences in the first four positions (**E**D**LA** for spSrtA versus **T**D**IE** for spySrtA, differences in **bold**). Based on our previous work, we attribute the lower relative activities for the G-, S-, and A-containing peptides in spySrtA_pneumoniae_ to the β7-β8^+1^ Glu.^12^ Specifically, we predicted this may be due to an interaction with the β6^-2^ R184 residue in spSrtA, which was supported by the observation that E208A and E208G, both at the β7-β8^+1^ position, spSrtA mutants each revealed ≥ 2-fold increases in relative reaction rates for all three peptides.^12^ Therefore, we *in silico* created T209E in the wild-type spySrtA structure to probe this hypothesis, and indeed saw that different rotamers of the mutated Glu are within distances consistent with forming a non-covalent interaction with the β6^-2^ T185 from the spySrtA scaffold (**Figure S2b**). Taken together, we consider this interaction to be a likely cause for the reduced activity in spySrtA_pneumoniae_.

### Structure determination and analysis of wild-type sagSrtA_247_ protein

Our rationale for choosing to investigate the role of β7-β8 loop residues in *S. pyogenes* and *S. agalactiae* was to develop a structural model for probing these selectivity determinants. Both spySrtA and sagSrtA_238_ were previously crystallized and we reasoned that experimental structural data would enable an understanding of the stereochemistry of sortase-substrate interactions in a way not available to spSrtA, which crystallizes as a catalytically inactive domain-swapped dimer.^40,41,43,45^

Beginning with sagSrtA, an important consideration for the reported sagSrtA_238_ structure was that we found this variant to be inactive in our enzyme assays (**Figure 2b**). The crystal structure of sagSrtA_238_ shows a dodecameric protein comprised of two hexameric rings (**Figure S4a**). The asymmetric unit contains one-and-a-half of these units, or 18 protomers total, with the other half of the second dodecamer present in a molecule related by symmetry. Each protomer is bound to three zinc ions, with additional ions modeled in the overall structure. There is no known biological requirement for higher order oligomers in sagSrtA activity or for zinc-binding and presumably, the presence of these ions is due to the crystallization conditions, e.g., zinc acetate or zinc sulfate.^44^ This previous work was focused on comparing the structures of sagSrtA_238_ with the *S. agalactiae* Class C1 sortase and did not include enzyme activity data.^45^ Of particular interest to us, however, is that in all 18 protomers of the sagSrtA_238_ asymmetric unit, there are unresolved residues in either the β4-β5 loop, β7-β8 loop, or both (**Figure S4b**). Based on our enzyme assay data and previous structural analyses, we therefore sought to crystallize and determine a structure of sagSrtA_247_ that would display relevant loop residues in an enzymatically active protein construct.^12^

Crystallization conditions for sagSrtA_247_ were identified using the Hampton PEGR_X_ screen and optimized, as described in the Materials and Methods. We ultimately solved the structure of sagSrtA_247_ to *R*_work_/*R*_free_ = 0.186/0.207 at 1.4 Å resolution, which includes residues Q82-L247 (**Figure 4**). All diffraction and refinement statistics are in **Table 1**.

**Table 1.**
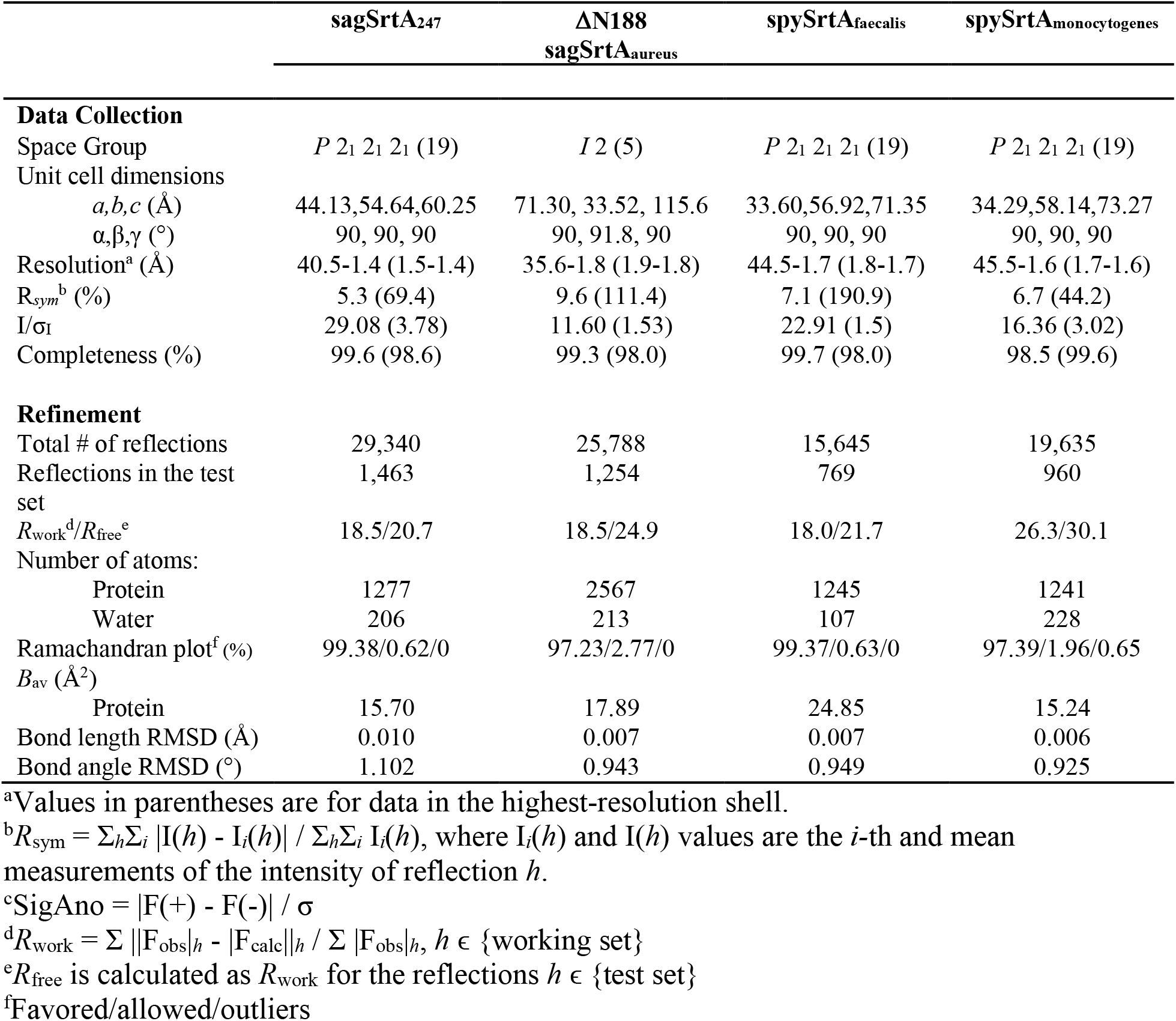
Data Collection and Refinement Statistics.

| | sagSrtA<sub>247</sub> | ΔN188<br>sagSrtA<sub>aureus</sub> | spySrtA<sub>faecalis</sub> | spySrtA<sub>monocytogenes</sub> |
| --- | --- | --- | --- | --- |
| **Data Collection** |  |  |  |  |
| Space Group | *P* 2<sub>1</sub> 2<sub>1</sub> 2<sub>1</sub> (19) | *I* 2 (5) | *P* 2<sub>1</sub> 2<sub>1</sub> 2<sub>1</sub> (19) | *P* 2<sub>1</sub> 2<sub>1</sub> 2<sub>1</sub> (19) |
| Unit cell dimensions |  |  |  |  |
| *a, b, c* (Å) | 44.13, 54.64, 60.25 | 71.30, 33.52, 115.6 | 33.60, 56.92, 71.35 | 34.29, 58.14, 73.27 |
| $\alpha, \beta, \gamma$ (°) | 90, 90, 90 | 90, 91.8, 90 | 90, 90, 90 | 90, 90, 90 |
| Resolution<sup>a</sup> (Å) | 40.5-1.4 (1.5-1.4) | 35.6-1.8 (1.9-1.8) | 44.5-1.7 (1.8-1.7) | 45.5-1.6 (1.7-1.6) |
| *R*<sub>sym</sub><sup>b</sup> (%) | 5.3 (69.4) | 9.6 (111.4) | 7.1 (190.9) | 6.7 (44.2) |
| *I*/σ<sub>*I*</sub> | 29.08 (3.78) | 11.60 (1.53) | 22.91 (1.5) | 16.36 (3.02) |
| Completeness (%) | 99.6 (98.6) | 99.3 (98.0) | 99.7 (98.0) | 98.5 (99.6) |
| **Refinement** |  |  |  |  |
| Total # of reflections | 29,340 | 25,788 | 15,645 | 19,635 |
| Reflections in the test set | 1,463 | 1,254 | 769 | 960 |
| *R*<sub>work</sub><sup>d</sup>/*R*<sub>free</sub><sup>e</sup> | 18.5/20.7 | 18.5/24.9 | 18.0/21.7 | 26.3/30.1 |
| Number of atoms: |  |  |  |  |
| Protein | 1277 | 2567 | 1245 | 1241 |
| Water | 206 | 213 | 107 | 228 |
| Ramachandran plot<sup>f</sup> (%) | 99.38/0.62/0 | 97.23/2.77/0 | 99.37/0.63/0 | 97.39/1.96/0.65 |
| *B*<sub>av</sub> (Å<sup>2</sup>) |  |  |  |  |
| Protein | 15.70 | 17.89 | 24.85 | 15.24 |
| Bond length RMSD (Å) | 0.010 | 0.007 | 0.007 | 0.006 |
| Bond angle RMSD (°) | 1.102 | 0.943 | 0.949 | 0.925 |

<sup>a</sup>Values in parentheses are for data in the highest-resolution shell.
<sup>b</sup> $R_{\text{sym}} = \sum_h \sum_i |I(h) - I_i(h)| / \sum_h \sum_i I_i(h)$ , where $I_i(h)$ and $I(h)$ values are the *i*-th and mean measurements of the intensity of reflection *h*.
<sup>c</sup> $\text{SigAno} = |F(+)-F(-)| / \sigma$
<sup>d</sup> $R_{\text{work}} = \sum ||F_{\text{obs}}|_h - |F_{\text{calc}}||_h / \sum |F_{\text{obs}}|_h$ , $h \in \{\text{working set}\}$
<sup>e</sup> $R_{\text{free}}$ is calculated as $R_{\text{work}}$ for the reflections $h \in \{\text{test set}\}$
<sup>f</sup>Favored/allowed/outliers

**Figure 4.**
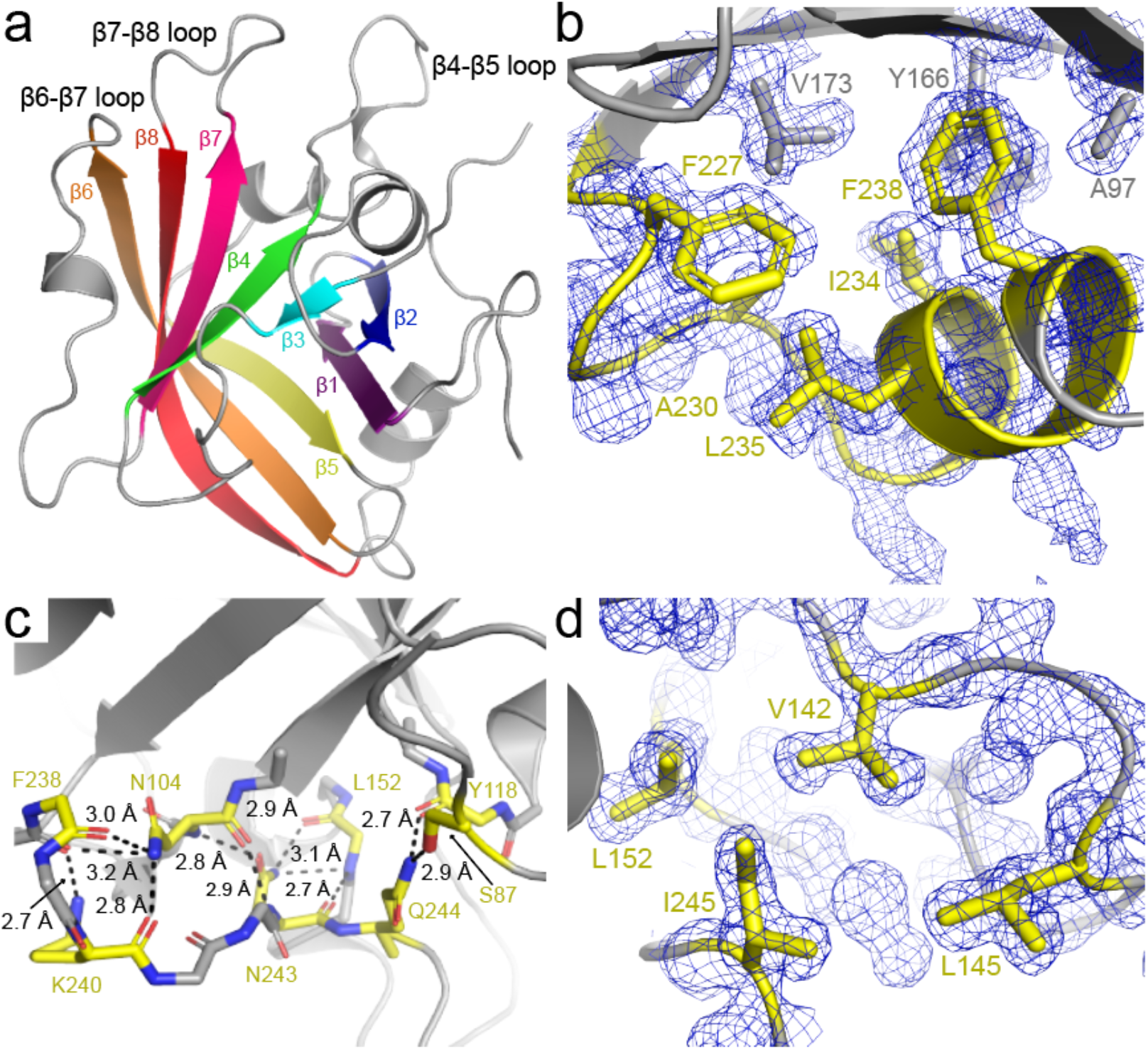
Structural characteristics of sagSrtA_247_. (**a**) SagSrtA_247_ adopts the conserved sortase fold. The protein is in gray cartoon, with β-strands numbered and colored as labeled. (b) C-terminal residues in sagSrtA_247_ that were in the construct previously crystallized, but were not resolved in the structure, make intra-protein hydrophobic interactions. Unresolved residues from PDB ID 3RCC are in yellow, with other sagSrtA_247_ residues in gray. Residues involved in the interaction are labeled and their side chains are shown as sticks. The electron density in this region is also shown, the 2*F*_o_-*F*_c_ map is rendered at 1σ. (**c**) C-terminal residues not included in the previously crystallized construct make several interactions in sagSrtA_247_. All distances are labeled and residues involved in contacts (including main chain of residues 238-244) are shown as sticks and colored by heteroatom (C=yellow, O=red, N=blue). Side chain atoms are shown when involved in the interaction, otherwise they are omitted for visual clarity. (**d**) The hydrophobic interaction involving I245 is shown, all residues are colored and the electron density is as in (**b**).

The sagSrtA_247_ structure adopts the conserved sortase fold, with a closed 8-stranded antiparallel β-sheet at its core (**Figure 4a**).^2^ Residues G225-F238, which are present in the crystallized sagSrtA_238_ construct, but are not resolved in that crystal structure, form a C-terminal helix that directly interacts with residues in the β1, β2, β5, and β6 strands, in a hydrophobic manner (**Figure 4b**). There are also several hydrogen bonds formed in residues C-terminal to F238, specifically S239, K240, N243, and Q244 which are not present in the crystallized sagSrtA_238_ construct. These interactions are largely mediated by mainchain atoms, but also include the sidechains of S87, N104, K240, N243, and Q244 (**Figure 4c**). In addition, the side chain of I245 is a part of a hydrophobic pocket formed with V142, L145, and L152, which are residues in the β4-β5 loop (**Figure 4d**). We predict that the lack of these interactions destabilizes the sagSrtA_238_ monomer, resulting in an inactive enzyme.

Interestingly, the C-terminus of saSrtA is substantially shorter than that of sagSrtA, or other *Streptococcus* SrtA proteins. Alignment of available saSrtA structures, including PDB IDs 1IJA (NMR), 1T2P (X-ray crystallography), and 2KID (NMR, +LPAT* peptidomimetic) indicate that the C-terminus of saSrtA, K206 (using 2KID and 1T2P numbering), corresponds stereochemically to K223 in sagSrtA (**Figure S5a**).^15,20,48^ Structural analyses of the two hydrophobic pockets that involve C-terminal residues in sagSrtA_247_ suggest that the saSrtA sequence would be unlikely to accommodate a similar C-terminal extension (**Figures S5b-c**). Specifically, an overlay of relevant structures suggests steric clashes between E77 in saSrtA with F238 in sagSrtA_247_ and R124 in saSrtA with I245 in sagSrtA_247_ (**Figures S5b-c**).

Structural alignments with protomers in sagSrtA_238_ (PDB ID 3RCC), spySrtA (3FN5), and *S. mutans* SrtA (4TQX) reveal that overall, sagSrtA_247_ adopts a conformation most similar to that of spySrtA (**Figure S6a**). Alignment with main chain atoms in each of the 18 protomers of the sagSrtA_238_ asymmetric unit reveal an average root-mean squared deviation (RMSD) value of 0.690 Å over 384 atoms, with the highest similarity between our structure and chain O (0.544 Å over 371 atoms) and lowest with chain K (0.792 Å over 394 atoms). Alignment with the two protomers of spySrtA revealed RMSD values of: 0.503 Å (551 atoms, with chain A) and 0.475 Å (535 atoms, chain B), and with *S. mutans* SrtA, the main-chain atoms align with an RMSD value of 0.566 Å over 549 atoms (**Figure S6a**). The largest differences between sagSrtA_247_ and spySrtA occur at the N-termini of both proteins (**Figure S6b**). In addition, we see ∼1 Å shifts in two of the structurally-conserved loops that surround the peptide-binding cleft, the β4-β5 and β7-β8 loops, likely due to differences in crystal packing (**Figure S6c**). Notably, the side-chain location and orientation of residues in these loops previously identified as being selectivity determinants of spSrtA activity are stereochemically conserved (**Figure S6d**).^12^ Taken together, our structure suggests that the active sagSrtA protein adopts a similar monomeric conformation as spySrtA.

### Structural analyses of chimeric Streptococcus SrtA proteins

We next wanted to investigate how our β7-β8 loop chimeras affect the structures of sagSrtA and spySrtA. We attempted to crystallize all 8 of our loop chimeras, using previously optimized conditions for sagSrtA and spySrtA, as well as by setting up commercially available crystal screens (e.g., Hampton PEG/ION, Index, and/or PEGRx). We were able to crystallize and solve structures of two of our chimeric proteins: spySrtA_faecalis_ and spySrtA_monocytogenes_ (**Table 1**). In addition, we successfully crystallized spySrtA_pneumoniae_ and sagSrtA_pneumoniae_, but they were not of diffraction quality.

We resolved all residues of the β7-β8 loop of our spySrtA_faecalis_ structure (**Figure S7a**), and all but the middle two residues of the spySrtA_monocytogenes_ loop (**Figure S7b**). The overall spySrtA variant conformations are identical to the wild-type protein, and alignments of mainchain atoms revealed RMSD values of 0.351 Å (533 atoms) and 0.252 Å (488 atoms) for spySrtA_faecalis_ and spySrtA_monocytogenes_, respectively (**Figure S7c**).

In the spySrtA variant structures, the orientation of the spySrtA_faecalis_ loop is very similar to the wild-type protein (**Figure 5a**), and the intra-loop hydrogen bond between the conserved β7-β8^+2^ Asp and β7-β8^+6^ Thr is maintained. This is not the case in the spySrtA_monocytogenes_ β7-β8 loop, as compared to the *L. monocytogenes* SrtA (lmSrtA) structure (**Figures 5b-c**).^49^ Here, the wild-type position of the β7-β8^+3^ Pro sterically clashes with the β4-β5^+3^ F145 residue in spySrtA and as a result, the β7-β8^+3^ Pro in spySrtA_monocytogenes_ is shifted away relative to the β4-β5 loop (**Figures 5b-c**). This results in breakage of the intra-loop hydrogen bond; whereas the distance between β7-β8^+1^ D118 and β7-β8^+6^ T123 is 2.8 Å in lmSrtA, it is 7.7 Å in spySrtA_monocytogenes_ (black arrows in **Figures 5b-c**).

**Figure 5.**
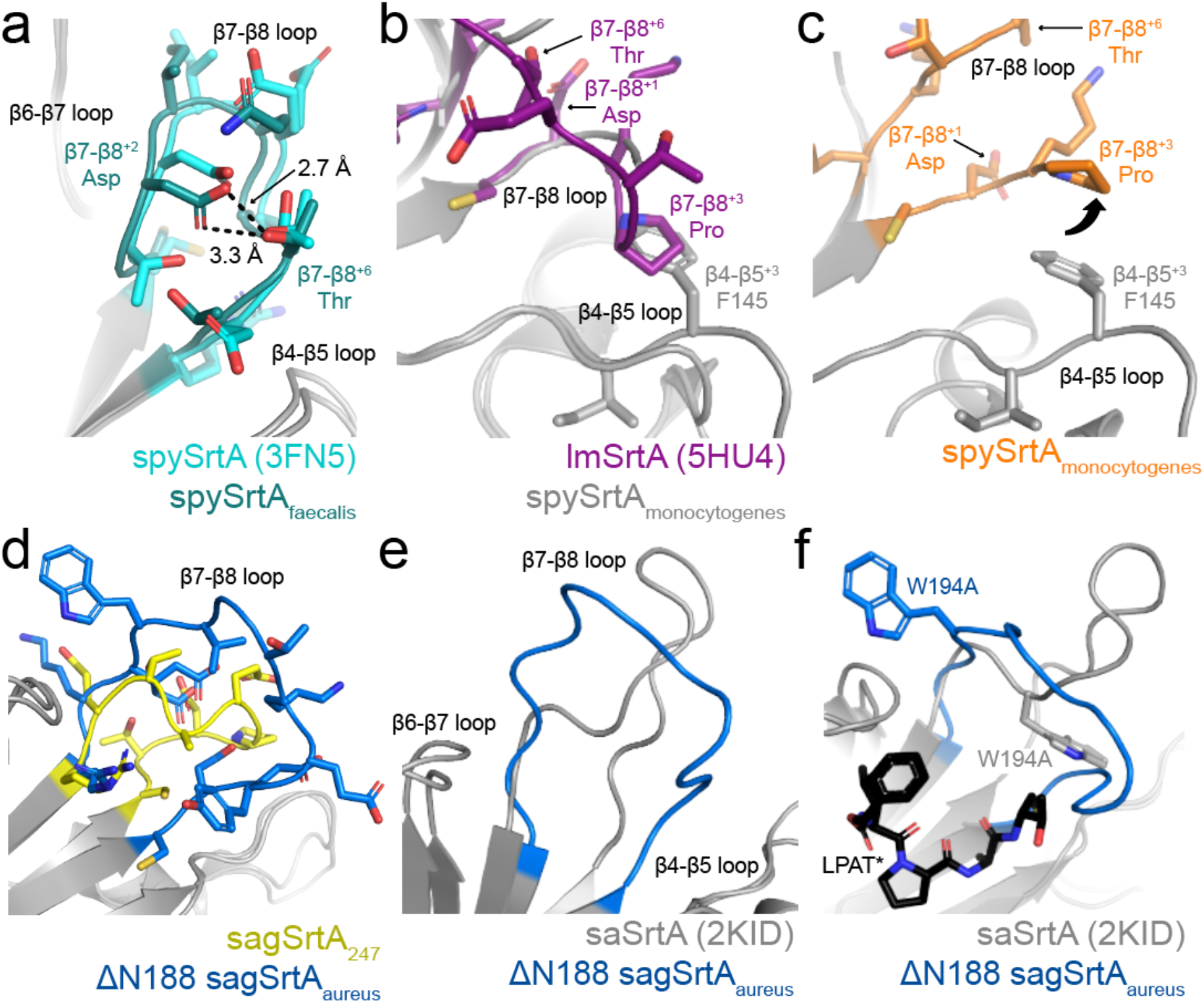
Structural characteristics of sagSrtA and spySrtA β7-β8 loop chimeras. In all, relevant residues are colored by structure as labeled and when represented, side chains are shown as sticks and colored by heteroatom (O=red, N=blue). Other residues are shown as gray cartoon. The intraloop hydrogen bond is conserved between spySrtA (PDB ID 3FN5) and spySrtA_faecalis_. The β7-β8^+3^ Pro in lmSrtA (5HU4) occupies the same position as the β4-β5^+3^ Phe of spySrtA_monocytogenes_, suggesting why the β7-β8 loop in spySrtA_monocytogenes_ is not well ordered and this variant is not as active. Furthermore, the intraloop hydrogen bond is not maintained in spySrtA_monocytogenes_, as compared to lmSrtA. The black arrows indicate the residues involved in this interaction in lmSrtA. (**c**) The β7-β8^+3^ Pro in spySrtA_monocytogenes_ is translated up, as compared to that of lmSrtA in (**b**). The black arrows show the relevant residues for the intra-loop hydrogen bond that is not conserved, as compared to (**b**). (**d-f**) Comparison of the ΔN188 sagSrtA_aureus_ β7-β8 loop with sagSrtA_247_ (**d**) and saSrtA (2KID, **e-f**). In (**f**), the LPAT* peptidomimetic is shown as black sticks and colored by heteroatom. The different positions of W194 are labeled.

In the active conformation of saSrtA (PDB ID 2KID), the 12 residues in the β7-β8 loop adopt a tight structure, mediated by several intraloop hydrogen bonds and noncovalent interactions, all of which include the sidechain atoms of N188 (**Figures 5e, S7d**). Therefore, we wanted to test the contribution of this residue on the spySrtA_aureus_ and sagSrtA_aureus_ proteins, with the variants ΔN188 spySrtA_aureus_ and ΔN188 sagSrtA_aureus_ (**Figures 3a-b**). These proteins were similar to other saSrtA loop variants in that they were selective for LPAT**G**; however, the relative activities were reduced by half as compared to spySrtA_aureus_ and 4.5-fold as compared to sagSrtA_aureus_, respectively (**Figures 3a-b**). We next crystallized and solved the structure of ΔN188 sagSrtA_aureus_, and were able to resolve all 11 residues in the loop in one of the protomers (**Figure S7e**). Alignment of the sagSrtA_247_ and ΔN188 sagSrtA_aureus_ structures reveals the greatest structural variability in the β6-β7 and β7-β8 loops, and the overall RMSD = 0.298 Å (496 mainchain atoms) (**Figure S7f**).

Comparison of the β7-β8 loops in ΔN188 sagSrtA_aureus_ and sagSrtA_247_ indicate that the loop in ΔN188 sagSrtA_aureus_ adopts a more *open* shape (**Figure 5d**). In the absence of N188, it is unsurprising that the β7-β8 loop in ΔN188 sagSrtA_aureus_ is missing the equivalent saSrtA interactions and shows only very weak, and likely unfavorable repulsive electrostatic ones between D185 and E195 (**Figure S7g**). Finally, we see displacement of the W194 residue in the ΔN188 sagSrtA_aureus_ structure (**Figure 5f**). We hypothesize this is the largest contributor to the weaker activity in ΔN188 sagSrtA_aureus_, as mutation to alanine at this residue was previously shown to reduce the activity of saSrtA by approximately 2-fold, which is of a similar magnitude to the effect of ΔN188 on spySrtA_aureus_ (**Figure 3a-b**).^14^ Interestingly, despite differences in overall shape of the β7-β8 loop in ΔN188 sagSrtA_aureus_, the protein retains the stringent selectivity of the saSrtA protein, recognizing only LPAT**G** (**Figure 3b**).

### Mutagenesis of Streptococcus SrtA proteins

Based on our structures, it is not immediately clear why sagSrtA_247_ is less active than spySrtA (**Figures 3a-b**). It is also not obvious why, for example, the ΔN188 mutation described above reduced spySrtA_aureus_ activity 2-fold, but sagSrtA_aureus_ activity 4.5-fold, or why there appears to be reduced ability to recognize S-containing peptides for several of the sagSrtA variants, as compared to spySrtA (**Figures 3a-b**). In the vicinity of the peptide-binding cleft, there are 4 non-conservative mutations (**Figure 6a**). Of these, we identified two that may contribute directly to the relatively low activity of sagSrtA_247_: K183 and P209.

**Figure 6.**
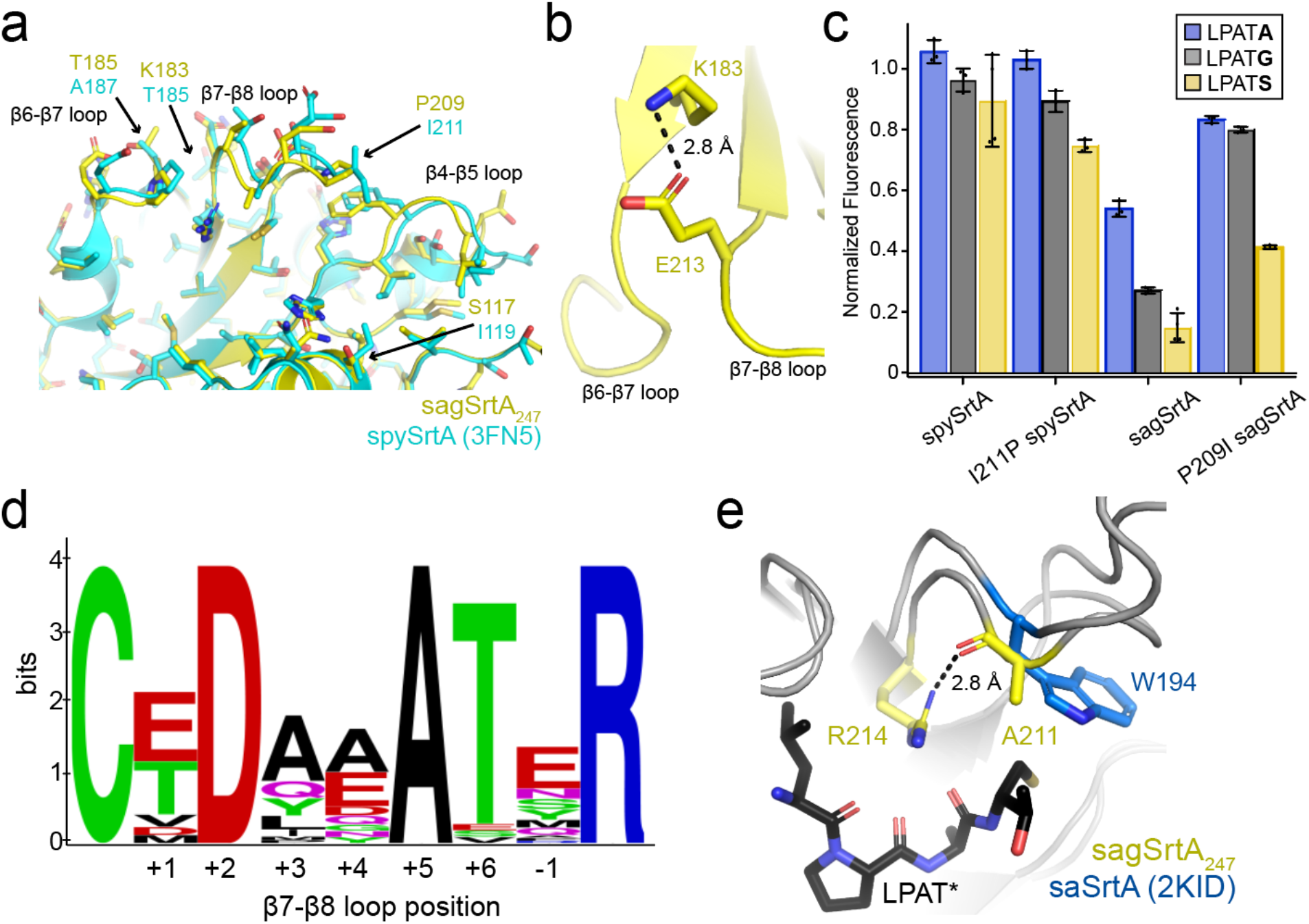
Residues in the β7-β8 loop of *Streptococcus* proteins that regulate enzyme function. (**a**) Although the peptide-binding pockets of sagSrtA and spySrtA are well conserved, there are four non-conservative amino acid differences, as labeled. Both proteins are shown in cartoon representation, with side chains as sticks and colored by heteroatom. The β4-β5, β6-β7, and β7-β8 loops are also labeled. (**b**) The side chain of β7-β8^-1^ E213 interacts with that of β6^-2^ K183 in sagSrtA_247_. This is an interaction that negatively affects enzyme activity for spSrtA, as previously reported, and which spySrtA does not share.^12^ (**c**) Mutation of the β7-β8^+3^ residue in sagSrtA from Pro to Ile, as in spySrtA, increases activity. However, the converse mutation in spySrtA, from Ile to Pro, has little to no effect. Assays were run and data was collected as in **Figures 2-3**. Averaged assay values and standard deviations are in **Table S1**. (**d**) WebLogo analysis of 37 *Streptococcus* SrtA proteins from the UniProt database. All sequences are in **Table S2**. (**e**) The absolutely conserved β7-β8^+5^ Ala in the sequences in (**d**) is stereochemically located in the same position as the W194 residue in saSrtA (2KID). A hydrogen bond between the A211 carbonyl and guanidinium group of the catalytic R214 is labeled.

Beginning with the K183 residue of sagSrtA_247_, we noted that it occupied the β6^-2^ position of the enzyme. In spySrtA, a threonine (T185) is present at the β6^-2^ position, which should not interact with the β7-β8^-1^ Glu, thereby avoiding an interaction that was previously shown to reduce enzyme activity in spSrtA.^12^ The Lys substitution in sagSrtA, however, would allow the β6^-2^ K183 to interact with the β7-β8^-1^ E213 (**Fig 6b**), and potentially reduce activity in a manner similar to the hypothesized interaction of the β6^-2^ R184 with β7-β8^-1^ E214 in spSrtA.^12^

With respect to the second residue (P209), we noticed that it occupied the β7-β8^+3^ loop position in sagSrtA_247_, similar to that in lmSrtA. We previously hypothesized that the β7-β8^+3^ Pro negatively affected spSrtA_monocytogenes_ and mutation of the wild-type Leu in L209P spSrtA reduced activity by about 2-fold.^12^ To test this, we mutated the β7-β8^+3^ loop residue in sagSrtA_247_ and spySrtA to that of the other protein, or P209I sagSrtA_247_ and I211P spySrtA, respectively. Relative enzyme activities were assayed and indeed, we saw an ∼2-fold increase in activity for P209I sagSrtA for G-, S-, and A-containing peptides (**Figure 6c**). Interestingly, we saw minimal differences in the activities of I211P spySrtA as compared to wild-type spySrtA, suggesting that the β7-β8^+3^ residue interaction may not be as critical for spySrtA, which is relatively more active than either sagSrtA or spSrtA (**Figures 2b**).

### Sequence Patterns in the β7-β8 loops of Streptococcus SrtA proteins

Finally, we wanted to gain a general understanding of sequence patterns in the β7-β8 loops of *Streptococcus* SrtA proteins. Therefore, we created a WebLogo of 37 β7-β8 loops from *Streptococcus* SrtA proteins from the UniProt database (**Figure 6d, Table S2**).^50,51^ We identified the loop sequences by using the catalytic cysteine and arginine residues to mark the N- and C-terminal residues of the β7-β8 loop, respectively. Our WebLogo analysis agreed with our biochemical and structural observations. The β7-β8^+2^ residue is an Asp in all of the loops, while the β7-β8^+6^ residue is a Thr (or Ser) in 35/37 sequences, consistent with an intraloop hydrogen bond interaction observed here and previously.^12^ We also observed an interaction between the β7-β8^+3^ position and the β4-β5 loop, typically of a hydrophobic nature.^12^ Consistent with this, our WebLogo analysis showed that the β7-β8^+3^ position is hydrophobic (Ala, Ile, Leu, Met, or Tyr) in 31 of the sequences, with the remaining sequences containing either Gln (5/37 sequences) or a Pro that is present in only the sagSrtA enzyme. It is unclear if the Gln can interact with the β4-β5 residues previously identified and if, as discussed for spySrtA, this interaction is correlated to the presence of a β7-β8^-1^/β6^-2^ interaction.

Notably, all 37 sequences contain a β7-β8^+5^ Ala residue (**Table S2**). Analyses of this residue in the sagSrtA_247_ and spySrtA structures show that it is solvent exposed (**Figure S8**). However, alignment of sagSrtA_247_ with saSrtA-LPAT* (2KID) revealed that the β7-β8^+5^ A211 sidechain points directly towards the peptide (**Figure 6e**). Furthermore, the carbonyl of A211 interacts with the guanidinium group of the catalytic arginine and the Ala is in the same stereochemical position as W194 in saSrtA (**Figure 6e**). Taken together, this suggests that the β7-β8^+5^ Ala residue in *Streptococcus* SrtA proteins may play an important role in enzyme function, just as the β7-β8^+10^ Trp residue does in saSrtA.

## Discussion

Work from ourselves and others indicates that the structurally conserved, yet sequence variable, β6-β7 and β7-β8 loops in Class A sortases directly affect target recognition and enzyme activity.^12,38,39^ Previously, we used the Class A sortase from *Streptococcus pneumoniae* to investigate the differences in selectivity and activity at the P1’ position as compared to SrtA from *Staphylococcus aureus* and 7 other organisms.^11,12^ Here, we extended these studies to look at similar chimeric SrtA enzymes from *Streptococcus pyogenes* and *Streptococcus agalactiae*, which were previously crystallized.^43–45^ Using protein biochemistry and structural biology, we find additional evidence in support of our hypothesis that the β7-β8 loop residues in these proteins determine overall enzyme activity and selectivity in a similar manner to spSrtA. Specifically, our data strongly supports the presence of 3 interactions mediated by β7-β8 loop residues in *Streptococcus* SrtA proteins that can mediate enzyme function.^12^ Although the exact nature of these interactions can vary in SrtA proteins from different organisms, e.g., *Staphylococcus aureus*, we argue that related ones are likely present across the broad sortase superfamily.

Our work also highlights the need for additional sortase structures that are paired with biochemical data. For example, we discovered that the sagSrtA construct previously crystallized is of an inactive enzyme, and omits several important C-terminal interactions.^44,45^ This is also notable because the C-terminus of saSrtA is substantially shorter than that of the *Streptococcus* SrtA proteins and without the biochemical knowledge of enzyme activity, fundamental information about these enzymes is missed. In addition, the only available spSrtA structures in the Protein Data Bank are of domain-swapped dimers, which are not enzymatically active in our hands (data not shown).^40,41^ Considering the observations about contributions of individual residues to activity and/or selectivity by ourselves and others, there remains much to be learned from the study of individual sortase enzymes.

Finally, the work presented here may have implications for the continued development of sortase mediated-ligation as a tool for protein engineering. Recent applications of *sortagging* in cells and the evolution of sortases to recognize specific targets for potential therapeutics are amongst a number of exciting developments in the field.^6,52^ A greater understanding of substrate selectivity and target recognition could enable more sophisticated orthogonal labeling schemes in which multiple sortase enzymes can be utilized to recognize and modify distinct sequences on a single protein or simultaneous labeling of multiple targets.^3,9,53,54^ This ability to add numerous site-specific tags to protein targets *in vitro* and *in vivo* would be a powerful addition to the arsenal of protein engineering.

## Experimental Procedures

### Sequences used

The wild-type spySrtA sequence used is from the published structure, PDB ID 3FN5. This sequence was originally amplified from serotype M1 *S. pyogenes* strain SF370 genomic DNA, as previously described.^43^ This sequence is 74% identical (85% similar) to the *S. pyogenes* Class A sortase in UniProt, A0A2W5CEK0_STRPY (unreviewed). The wild-type sagSrtA sequence used is from the published structure, PDB ID 3RCC. This sequence was originally amplified from genomic DNA of *S. agalactiae* strain 2603V/R (locus tag of SAG0961), as previously described.^44^ This sequence is 99% identical, differing only at Q132, which is a proline in 3RCC, to *S. agalactiae* SrtA in UniProt, SRTA_STRA3 (reviewed). This substitution occurs in the β3-β4 loop. All constructs in this work, including chimeric and mutant proteins, were purchased from Genscript in the pET28a(+) vector.

### Protein expression and purification

All proteins were expressed and purified as previously described for related SrtA proteins.^11,12^ Briefly, plasmids were transformed into *Escherichia coli* BL21 (DE3) competent cells and grown in LB media, with protein induction at OD_600_ 0.6-0.8 using 0.15 mM IPTG for 18-20 h at 18 °C. The cells were harvested in lysis buffer [0.05 M Tris pH 7.5, 0.15 M NaCl, 0.5 mM ethylenediaminetetraacetic acid (EDTA)] and whole cell lysate was clarified using centrifugation. The supernatant was filtered and loaded onto a 5 mL HisTrap HP column (GE Life Sciences, now Cytiva), followed by washing [0.05 M Tris pH 7.5, 0.15 M NaCl, 0.02 M imidazole, 0.001 M TCEP] and then elution [wash buffer with 0.3 M imidazole] of the desired protein. The His-tags of proteins prepared for crystallography were proteolyzed using Tobacco Etch Virus (TEV) protease overnight at 4 °C and a ratio of ∼1:100 (TEV:protein). The His_6_-TEV sequence was left on proteins used for enzyme assays. Size exclusion chromatography (SEC) was conducted using a HiLoad 16/600 Superdex 75 column (GE Life Sciences, now Cytiva) in SEC running buffer [0.05 M Tris pH 7.5, 0.15 M NaCl, 0.001 M TCEP]. Purified protein corresponding to the monomeric peak was concentrated using an Amicon Ultra-15 Centrifugal Filter Unit (10,000 NWML) and analyzed by SDS-PAGE and analytical SEC.^12^ Protein concentrations were determined using theoretical extinction coefficients calculated using ExPASy ProtParam.^55^ Protein not immediately used was flash frozen in SEC running buffer and stored at -80°C.

### Crystallization

Prior to crystallization, spySrtA variants were dialyzed into crystallization buffer [20 mM Tris pH 7.5, 150 mM NaCl], based on previously published conditions.^43^ The protein concentrations used for crystallization were as follows: sagSrtA_247_ (15 mg/mL), ΔN188 sagSrtA_aureus_ (16 mg/mL), spySrtA_faecalis_ (42 mg/mL), and spySrtA_monocytogenes_ (20 mg/mL). The proteins were crystallized using the hanging drop vapor diffusion technique with well and protein solution mixed in a 1:1 ratio (2 μl:2 μl). Crystallization conditions for the spySrtA variants were optimized using the crystal conditions for the apo protein.^43^ For sagSrtA_247_, initial crystallization conditions were identified using the PEGRx screen from Hampton Research. The crystallization conditions of the crystals used for data collection were: sagSrtA_247_ [20% (*v/v*) 2-Propanol, 0.1 M MES monohydrate pH 6.1, 20% (*w/v*) PEG monomethyl ether 2,000], ΔN188 sagSrtA_aureus_ [12% (*v/v*) 2-Propanol, 0.02 M MES monohydrate pH 6, 24% (*w/v*) PEG monomethyl ether 2,000], spySrtA_faecalis_ [0.2 M sodium acetate, 0.1 M Tris pH 6, 30% (*w/v*) PEG 8,000], and spySrtA_monocytogenes_ [0.2 M sodium acetate, 0.1 M Tris pH 6.5, 24% (*w/v*) PEG 8,000]. For all proteins, glycerol was used as a cryoprotectant and the cryo solutions were equal to crystallization conditions plus 20% (*v/v*) glycerol for all except sagSrtA_247_ (plus 15% (*v/v*) glycerol). The crystals were flash-cooled by plunging into liquid nitrogen.

### Data collection, structure determination, and protein analyses

Initial data for sagSrtA_247_ were collected to 2.0 Å on a Bruker Apex CCD diffractometer at λ = 1.54056 nm. Data were collected at the Advanced Light Source (ALS) at Lawrence Berkeley National Laboratory (LBNL) on beamline 5.0.1 and 5.0.2, at λ= 1.00004 nm or 0.99988 nm over 360°, with Δϕ=0.25° frames and an exposure time of 0.5 s per frame. Data were processed using the XDS package (**Table 1**).^56,57^ Molecular Replacement was performed using Phenix with the following search models: spySrtA (PDB ID 3FN5) for spySrtA_faecalis_ and spySrtA_monocytogenes_, sagSrtA_238_ (3RCC) for sagSrtA_247_ and sagSrtA_247_ for ΔN188 sagSrtA_aureus_. Refinement was performed using Phenix, manual refinement was done using Coot, and model geometry was assessed using MolProbity and the PDB validation server.^58–60^ Phenix.Xtriage was also used to assess data quality, specifically to identify a number of outliers in the spySrtA_monocytogenes_ data.^60^ All crystal data and refinement statistics are in **Table 1**. Sequence alignments were performed using T-coffee or BlastP.^61,62^ Visualization of alignments were done using Jalview or Boxshade.^63^ WebLogo was also used to visualize sequences.^64^ Structural analyses and figure rendering were done using PyMOL. PDB accession codes for the structures presented here are: sagSrtA_247_ (7S56), ΔN188 sagSrtA_aureus_ (7S54), spySrtA_faecalis_ (7S57), and spySrtA_monocytogenes_ (7S53).

### Peptide synthesis

Model peptide substrates were synthesized via manual Fmoc solid phase peptide synthesis (SPPS) as previously described.^12^

### Fluorescence Assay for Sortas Activity

Enzyme assays were conducted using a Biotek Synergy H1 plate reader as previously described.^12^ The fluorescence intensity of each well was measured at 2-min time intervals over a 2-hr period at room temperature (λ_ex_ = 320 nm, λ_em_ = 420 nm, and detector gain = 75). All reactions were performed in at least triplicate. For each substrate sequence, the background fluorescence of the intact peptide in the absence of enzyme was subtracted from the observed experimental data. Background-corrected fluorescence data was then normalized to the fluorescence intensity of a benchmark reaction between wild-type saSrtA and Abz-LPATGG-K(Dnp).^12^ Data figures were prepared using GraphPad Prism 9.1.2.

## Acknowledgements

The authors would like to thank the other members of the Amacher and Antos labs for helpful discussions and assistance. They would also like to thank the Berkeley Center for Structural Biology (BCSB) for being an excellent resource for the crystallography community. The BCSB is supported in part by the National Institutes of Health, National Institute of General Medical Sciences, and the Howard Hughes Medical Institute. The Advanced Light Source is supported by the Director, Office of Science, Office of Basic Energy Sciences, of the U.S. Department of Energy under Contract No. DE-AC02-05CH11231. Other grant information: JFA and JMA were both funded by Cottrell Scholar Awards from the Research Corporation for Science Advancement. JFA was also funded by NSF CHE-CAREER-2044958. The Rigaku X-ray Diffractometer was funded by NSF CHE-MRI-1429164 and used to collect initial sagSrtA_247_ diffraction data. In addition, IMP and HMK received Elwha Undergraduate Summer Research Awards and DAJ received a Joseph & Karen Morse Student Research in Chemistry Fellowship to fund summer research.

## References

1. Spirig T, Weiner EM, Clubb RT (2011) Sortase enzymes in Gram-positive bacteria. Mol. Microbiol. 82:1044–1059.

2. Jacobitz AW, Kattke MD, Wereszczynski J, Clubb RT (2017) Sortase transpeptidases: structural biology and catalytic mechanism. Adv. Protein Chem. Struct. Biol. 109:223–264.

3. Antos JM, Truttmann MC, Ploegh HL (2016) Recent advances in sortase-catalyzed ligation methodology. Curr. Opin. Struct. Biol. 38:111–118.

4. Bradshaw WJ, Davies AH, Chambers CJ, Roberts AK, Shone CC, Acharya KR (2015) Molecular features of the sortase enzyme family. FEBS J. 282:2097–2114.

5. Dorr BM, Ham HO, An C, Chaikof EL, Liu DR (2014) Reprogramming the specificity of sortase enzymes. Proc. Natl. Acad. Sci. USA 111:13343–13348.

6. Podracky CJ, An C, DeSousa A, Dorr BM, Walsh DM, Liu DR (2021) Laboratory evolution of a sortase enzyme that modifies amyloid-β protein. Nat. Chem. Biol. 17:317–325.

7. Chen I, Dorr BM, Liu DR (2011) A general strategy for the evolution of bond-forming enzymes using yeast display. Proc. Natl. Acad. Sci. USA 108:11399–11404.

8. Piotukh K, Geltinger B, Heinrich N, Gerth F, Beyermann M, Freund C, Schwarzer D (2011) Directed evolution of sortase A mutants with altered substrate selectivity profiles. J. Am. Chem. Soc. 133:17536–17539.

9. Antos JM, Chew G-L, Guimaraes CP, Yoder NC, Grotenbreg GM, Popp MW-L, Ploegh HL (2009) Site-specific N-and C-terminal labeling of a single polypeptide using sortases of different specificity. J. Am. Chem. Soc. 131:10800–10801.

10. Schmohl L, Bierlmeier J, Gerth F, Freund C, Schwarzer D (2017) Engineering sortase A by screening a second-generation library using phage display. J Pept Sci 23:631–635.

11. Nikghalb KD, Horvath NM, Prelesnik JL, Banks OGB, Filipov PA, Row RD, Roark TJ, Antos JM (2018) Expanding the Scope of Sortase-Mediated Ligations by Using Sortase Homologues. Chembiochem 19:185–195.

12. Piper IM, Struyvenberg SA, Valgardson JD, Alex Johnson D, Gao M, Johnston K, Svendsen JE, Kodama HM, Hvorecny KL, Antos JM, et al. (2021) Sequence variation in the β7-β8 loop of bacterial Class A sortase enzymes alters substrate selectivity. J. Bio. Chem.:100981.

13. Malik A, Kim SB (2019) A comprehensive in silico analysis of sortase superfamily. J. Microbiol. 57:431–443.

14. Ton-That H, Mazmanian SK, Alksne L, Schneewind O (2002) Anchoring of surface proteins to the cell wall of Staphylococcus aureus. Cysteine 184 and histidine 120 of sortase form a thiolate-imidazolium ion pair for catalysis. J. Biol. Chem. 277:7447–7452.

15. Suree N, Liew CK, Villareal VA, Thieu W, Fadeev EA, Clemens JJ, Jung ME, Clubb RT (2009) The structure of the Staphylococcus aureus sortase-substrate complex reveals how the universally conserved LPXTG sorting signal is recognized. J. Biol. Chem. 284:24465–24477.

16. Naik MT, Suree N, Ilangovan U, Liew CK, Thieu W, Campbell DO, Clemens JJ, Jung ME, Clubb RT (2006) Staphylococcus aureus Sortase A transpeptidase. Calcium promotes sorting signal binding by altering the mobility and structure of an active site loop. J. Biol. Chem. 281:1817–1826.

17. Kappel K, Wereszczynski J, Clubb RT, McCammon JA (2012) The binding mechanism, multiple binding modes, and allosteric regulation of Staphylococcus aureus Sortase A probed by molecular dynamics simulations. Protein Sci. 21:1858–1871.

18. Ugur I, Schatte M, Marion A, Glaser M, Boenitz-Dulat M, Antes I (2018) Ca2+ binding induced sequential allosteric activation of sortase A: An example for ion-triggered conformational selection. PLoS One 13:e0205057.

19. Tee W-V, Guarnera E, Berezovsky IN (2020) Disorder driven allosteric control of protein activity. Current Research in Structural Biology 2:191–203.

20. Ilangovan U, Ton-That H, Iwahara J, Schneewind O, Clubb RT (2001) Structure of sortase, the transpeptidase that anchors proteins to the cell wall of Staphylococcus aureus. Proc. Natl. Acad. Sci. USA 98:6056–6061.

21. Chan AH, Yi SW, Terwilliger AL, Maresso AW, Jung ME, Clubb RT (2015) Structure of the Bacillus anthracis Sortase A Enzyme Bound to Its Sorting Signal: A FLEXIBLE AMINO-TERMINAL APPENDAGE MODULATES SUBSTRATE ACCESS. J. Biol. Chem. 290:25461–25474.

22. Goettig P, Brandstetter H, Magdolen V (2019) Surface loops of trypsin-like serine proteases as determinants of function. Biochimie 166:52–76.

23. Hedstrom L, Szilagyi L, Rutter WJ (1992) Converting trypsin to chymotrypsin: the role of surface loops. Science 255:1249–1253.

24. Pirola L, Zvelebil MJ, Bulgarelli-Leva G, Van Obberghen E, Waterfield MD, Wymann MP (2001) Activation loop sequences confer substrate specificity to phosphoinositide 3-kinase alpha (PI3Kalpha). Functions of lipid kinase-deficient PI3Kalpha in signaling. J. Biol. Chem. 276:21544–21554.

25. Shah NH, Wang Q, Yan Q, Karandur D, Kadlecek TA, Fallahee IR, Russ WP, Ranganathan R, Weiss A, Kuriyan J (2016) An electrostatic selection mechanism controls sequential kinase signaling downstream of the T cell receptor. Elife 5:e20105.

26. Laham LE, Mukhopadhyay N, Roberts TM (2000) The activation loop in Lck regulates oncogenic potential by inhibiting basal kinase activity and restricting substrate specificity. Oncogene 19:3961–3970.

27. Kunz J, Wilson MP, Kisseleva M, Hurley JH, Majerus PW, Anderson RA (2000) The activation loop of phosphatidylinositol phosphate kinases determines signaling specificity. Mol. Cell 5:1–11.

28. Nolen B, Taylor S, Ghosh G (2004) Regulation of protein kinases; controlling activity through activation segment conformation. Mol. Cell 15:661–675.

29. Liu H, Huang H, Voss C, Kaneko T, Qin WT, Sidhu S, Li SS-C (2019) Surface loops in a single SH2 domain are capable of encoding the spectrum of specificity of the SH2 family. Mol. Cell Proteomics 18:372–382.

30. Kaneko T, Huang H, Zhao B, Li L, Liu H, Voss CK, Wu C, Schiller MR, Li SS-C (2010) Loops govern SH2 domain specificity by controlling access to binding pockets. Sci. Signal. 3:ra34.

31. Kazlauskas A, Schmotz C, Kesti T, Hepojoki J, Kleino I, Kaneko T, Li SSC, Saksela K (2016) Large-Scale Screening of Preferred Interactions of Human Src Homology-3 (SH3) Domains Using Native Target Proteins as Affinity Ligands. Mol. Cell Proteomics 15:3270–3281.

32. Songyang Z, Shoelson SE, McGlade J, Olivier P, Pawson T, Bustelo XR, Barbacid M, Sabe H, Hanafusa H, Yi T (1994) Specific motifs recognized by the SH2 domains of Csk, 3BP2, fps/fes, GRB-2, HCP, SHC, Syk, and Vav. Mol. Cell. Biol. 14:2777–2785.

33. Teyra J, Huang H, Jain S, Guan X, Dong A, Liu Y, Tempel W, Min J, Tong Y, Kim PM, et al. (2017) Comprehensive Analysis of the Human SH3 Domain Family Reveals a Wide Variety of Non-canonical Specificities. Structure 25:1598–1610.e3.

34. Li SS-C (2005) Specificity and versatility of SH3 and other proline-recognition domains: structural basis and implications for cellular signal transduction. Biochem. J. 390:641–653.

35. Sparks AB, Rider JE, Hoffman NG, Fowlkes DM, Quillam LA, Kay BK (1996) Distinct ligand preferences of Src homology 3 domains from Src, Yes, Abl, Cortactin, p53bp2, PLCgamma, Crk, and Grb2. Proc. Natl. Acad. Sci. USA 93:1540–1544.

36. Brown T, Brown N, Stollar EJ (2018) Most yeast SH3 domains bind peptide targets with high intrinsic specificity. PLoS One 13:e0193128.

37. Feng S, Kasahara C, Rickles RJ, Schreiber SL (1995) Specific interactions outside the proline-rich core of two classes of Src homology 3 ligands. Proc. Natl. Acad. Sci. USA 92:12408–12415.

38. Bentley ML, Gaweska H, Kielec JM, McCafferty DG (2007) Engineering the substrate specificity of Staphylococcus aureus Sortase A. The beta6/beta7 loop from SrtB confers NPQTN recognition to SrtA. J. Biol. Chem. 282:6571–6581.

39. Wójcik M, Szala K, van Merkerk R, Quax WJ, Boersma YL (2020) Engineering the specificity of Streptococcus pyogenes sortase A by loop grafting. Proteins 88:1394–1400.

40. Biswas T, Misra A, Das S, Yadav P, Ramakumar S, Roy R (2020) Interrogation of 3D-swapped structure and functional attributes of quintessential Sortase A from Streptococcus pneumoniae. Biochem. J.

41. Misra A, Biswas T, Das S, Marathe U, Sehgal D, Roy RP, Suryanarayanarao R (2011) Crystallization and preliminary X-ray diffraction studies of sortase A from Streptococcus pneumoniae. Acta Crystallogr. Sect. F, Struct. Biol. Cryst. Commun. 67:1195–1198.

42. Huang X, Aulabaugh A, Ding W, Kapoor B, Alksne L, Tabei K, Ellestad G (2003) Kinetic mechanism of Staphylococcus aureus sortase SrtA. Biochemistry 42:11307–11315.

43. Race PR, Bentley ML, Melvin JA, Crow A, Hughes RK, Smith WD, Sessions RB, Kehoe MA, McCafferty DG, Banfield MJ (2009) Crystal structure of Streptococcus pyogenes sortase A: implications for sortase mechanism. J. Biol. Chem. 284:6924–6933.

44. Khare B, Samal A, Vengadesan K, Rajashankar KR, Ma X, Huang IH, Ton-That H, Narayana SVL (2010) Preliminary crystallographic study of the Streptococcus agalactiae sortases, sortase A and sortase C1. Acta Crystallogr. Sect. F, Struct. Biol. Cryst. Commun. 66:1096–1100.

45. Khare B, Krishnan V, Rajashankar KR, I-Hsiu H, Xin M, Ton-That H, Narayana SV (2011) Structural differences between the Streptococcus agalactiae housekeeping and pilus-specific sortases: SrtA and SrtC1. PLoS One 6:e22995.

46. Schmohl L, Bierlmeier J, von Kügelgen N, Kurz L, Reis P, Barthels F, Mach P, Schutkowski M, Freund C, Schwarzer D (2017) Identification of sortase substrates by specificity profiling. Bioorg. Med. Chem. 25:5002–5007.

47. Kruger RG, Dostal P, McCafferty DG (2004) Development of a high-performance liquid chromatography assay and revision of kinetic parameters for the Staphylococcus aureus sortase transpeptidase SrtA. Anal. Biochem. 326:42–48.

48. Zong Y, Bice TW, Ton-That H, Schneewind O, Narayana SVL (2004) Crystal structures of Staphylococcus aureus sortase A and its substrate complex. J. Biol. Chem. 279:31383–31389.

49. Li H, Chen Y, Zhang B, Niu X, Song M, Luo Z, Lu G, Liu B, Zhao X, Wang J, et al. (2016) Inhibition of sortase A by chalcone prevents Listeria monocytogenes infection. Biochem. Pharmacol. 106:19–29.

50. UniProt Consortium (2012) Reorganizing the protein space at the Universal Protein Resource (UniProt). Nucleic Acids Res. 40:D71–5.

51. Magrane M, Consortium U (2011) UniProt Knowledgebase: a hub of integrated protein data. Database (Oxford) 2011:bar009.

52. Wu Q, Ploegh HL, Truttmann MC (2017) Hepta-Mutant Staphylococcus aureus Sortase A (SrtA7m) as a Tool for in Vivo Protein Labeling in Caenorhabditis elegans. ACS Chem. Biol. 12:664–673.

53. Hess GT, Guimaraes CP, Spooner E, Ploegh HL, Belcher AM (2013) Orthogonal labeling of M13 minor capsid proteins with DNA to self-assemble end-to-end multiphage structures. ACS Synth. Biol. 2:490–496.

54. Guimaraes CP, Witte MD, Theile CS, Bozkurt G, Kundrat L, Blom AEM, Ploegh HL (2013) Site-specific C-terminal and internal loop labeling of proteins using sortase-mediated reactions. Nat. Protoc. 8:1787–1799.

55. Wilkins MR, Gasteiger E, Bairoch A, Sanchez JC, Williams KL, Appel RD, Hochstrasser DF (1999) Protein identification and analysis tools in the ExPASy server. Methods Mol. Biol. 112:531–552.

56. Kabsch W (2010) XDS. Acta Crystallogr. Sect. D, Biol. Crystallogr. 66:125–132.

57. Kabsch W (2010) Integration, scaling, space-group assignment and post-refinement. Acta Crystallogr. Sect. D, Biol. Crystallogr. 66:133–144.

58. Chen VB, Arendall WB, Headd JJ, Keedy DA, Immormino RM, Kapral GJ, Murray LW, Richardson JS, Richardson DC (2010) MolProbity: all-atom structure validation for macromolecular crystallography. Acta Crystallogr. Sect. D, Biol. Crystallogr. 66:12–21.

59. Emsley P, Lohkamp B, Scott WG, Cowtan K (2010) Features and development of Coot. Acta Crystallogr. Sect. D, Biol. Crystallogr. 66:486–501.

60. Adams PD, Afonine PV, Bunkóczi G, Chen VB, Davis IW, Echols N, Headd JJ, Hung L-W, Kapral GJ, Grosse-Kunstleve RW, et al. (2010) PHENIX: a comprehensive Python-based system for macromolecular structure solution. Acta Crystallogr. Sect. D, Biol. Crystallogr. 66:213–221.

61. Notredame C, Higgins DG, Heringa J (2000) T-Coffee: A novel method for fast and accurate multiple sequence alignment. J. Mol. Biol. 302:205–217.

62. Altschul SF, Gish W, Miller W, Myers EW, Lipman DJ (1990) Basic local alignment search tool. J. Mol. Biol. 215:403–410.

63. Clamp M, Cuff J, Searle SM, Barton GJ (2004) The Jalview Java alignment editor. Bioinformatics 20:426–427.

64. Crooks GE, Hon G, Chandonia JM, Brenner SE (2004) WebLogo: a sequence logo generator. Genome Res. 14:1188–1190.

