## Supplemental Material for "Structural and biochemical analyses of selectivity determinants in chimeric *Streptococcus* Class A sortase enzymes"

by

Melody Gao, D. Alex Johnson, Isabel M. Piper, Hanna M. Kodama, Justin E. Svendsen, Elise Tahti, Brandon Vogel, John M. Antos, Jeanine F. Amacher

**Table of Contents**

**Figure S1.** Comparison of the sequences of six *Streptococcus* SrtA proteins. 2

**Table S1.** Enzyme assay data for all variants. 3

**Figure S2.** Interaction between the β7-β8 loop and $\beta6^{-2}$ residue in *Streptococcus* SrtA proteins. 4

**Figure S3.** Some chimeric proteins reveal differences in initial rates of the reaction. 5

**Figure S4.** The $sagSrtA_{238}$ structure. 6

**Figure S5.** Comparison of the C-termini of $sagSrtA_{247}$ and saSrtA. 7

**Figure S6.** The $sagSrtA_{247}$ structure is most similar to spySrtA (PDB ID 3FN5). 8

**Figure S7.** The β7-β8 loop sequences of *Streptococcus* proteins. 10

**Table S2.** The β7-β8 loop sequences of *Streptococcus* proteins. 11

**Figure S8.** A conserved alanine in the β7-β8 loops of *Streptococcus* proteins. 12

a

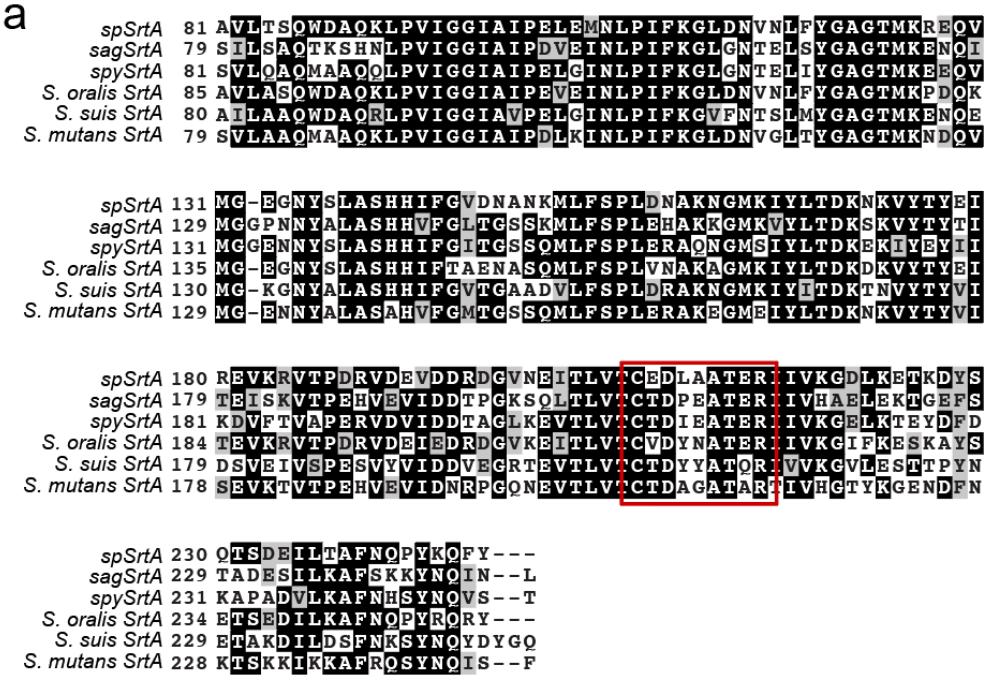


b

Pairwise sequence identity % (number of residues aligned)

| | spSrtA | sagSrtA | spySrtA | *S. oralis* SrtA | *S. suis* SrtA | *S. mutans* SrtA |
|---|---|---|---|---|---|---|
| spSrtA | 100% | 57% (167) | 63% (166) | 81% (167) | 62% (166) | 64% (166) |
| sagSrtA | - | 100% | 65% (168) | 58% (166) | 58% (168) | 64% (169) |
| spySrtA | - | - | 100% | 63% (166) | 63% (166) | 69% (168) |
| *S. oralis* SrtA | - | - | - | 100% | 61% (165) | 64% (165) |
| *S. suis* SrtA | - | - | - | - | 100% | 60% (168) |
| *S. mutans* SrtA | - | - | - | - | - | 100% |

**Figure S1. Comparison of the sequences of six *Streptococcus* SrtA proteins.** (**a**) The sequences of SrtA proteins from *S. pneumoniae* (spSrtA), *S. agalactiae* (sagSrtA), *S. pyogenes* (spySrtA), *S. oralis*, *S. mutans*, and *S. suis* were aligned using T-coffee and visualized with Boxshade. The β7-β8 loop residues are indicated with a red box. (**b**) Table of pairwise alignments of the six *Streptococcus* SrtA proteins reveal the closest sequence identity between spSrtA and *S. oralis* SrtA (81% identity) and least between spSrtA and sagSrtA (57% identity). Most values are in the 60-69% range (11 of the total 15).

**Table S1**. Enzyme assay data for all variants (N ≥ 3 for all).

| | Average normalized fluorescence at t = 2 h | | |
|---|---|---|---|
| | LPAT**X** | | |
| *S. pyogenes* SrtA variant | **G** | **S** | **A** |
| spySrtA | 0.97 ± 0.04 | 0.89 ± 0.15 | 1.05 ± 0.04 |
| $spySrtA_{aureus}$ | 0.97 ± 0.03 | 0.00 ± 0.00 | 0.05 ± 0.00 |
| $spySrtA_{faecalis}$ | 0.97 ± 0.06 | 0.97 ± 0.05 | 1.02 ± 0.05 |
| $spySrtA_{monocytogenes}$ | 0.45 ± 0.02 | 0.01 ± 0.00 | 0.06 ± 0.00 |
| $spySrtA_{pneumoniae}$ | 0.77 ± 0.04 | 0.72 ± 0.06 | 0.81 ± 0.03 |
| ΔN188 $spySrtA_{aureus}$ | 0.54 ± 0.02 | 0.01 ± 0.00 | 0.03 ± 0.00 |
| I215P spySrtA | 0.89 ± 0.04 | 0.75 ± 0.02 | 1.03 ± 0.03 |

| | Average normalized fluorescence at t = 2 h | | |
|---|---|---|---|
| | LPAT**X** | | |
| *S. agalactiae* SrtA variant | **G** | **S** | **A** |
| $sagSrtA_{247}$ | 0.27 ± 0.01 | 0.15 ± 0.05 | 0.54 ± 0.03 |
| $sagSrtA_{238}$ | 0.02 ± 0.00 | -0.02 ± 0.03 | 0.02 ± 0.00 |
| $sagSrtA_{aureus}$ | 0.90 ± 0.05 | -0.00 ± 0.00 | 0.02 ± 0.00 |
| $sagSrtA_{faecalis}$ | 0.67 ± 0.01 | 0.39 ± 0.03 | 0.66 ± 0.05 |
| $sagSrtA_{monocytogenes}$ | 0.08 ± 0.00 | -0.01 ± 0.00 | 0.00 ± 0.00 |
| $sagSrtA_{pneumoniae}$ | 0.25 ± 0.02 | 0.14 ± 0.01 | 0.21 ± 0.01 |
| ΔN188 $sagSrtA_{aureus}$ | 0.20 ± 0.06 | -0.01 ± 0.00 | -0.00 ± 0.00 |
| P209I sagSrtA | 0.80 ± 0.00 | 0.41 ± 0.00 | 0.83 ± 0.01 |

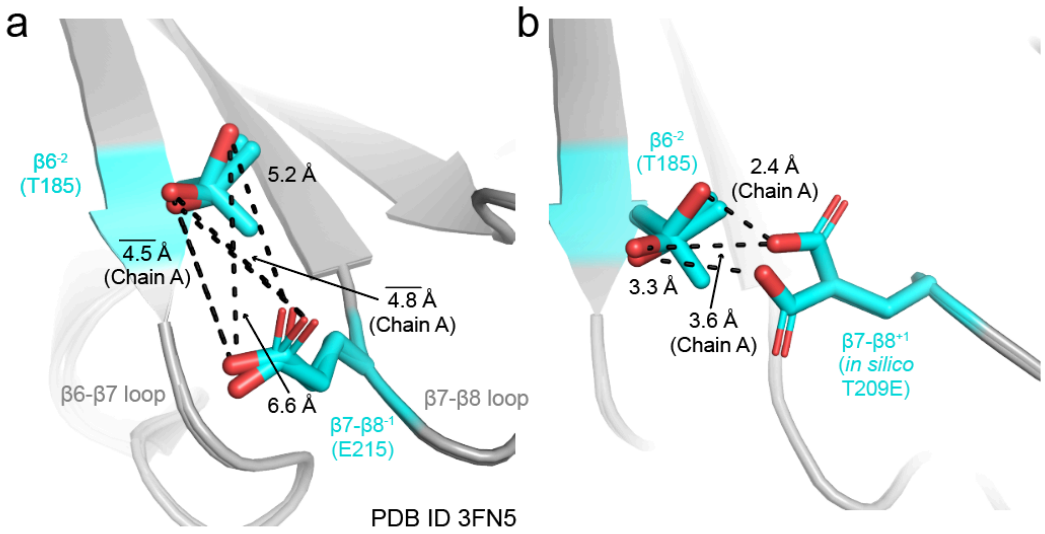


**Figure S2. Interaction between the β7-β8 loop and β6$^{-2}$ residue in *Streptococcus* SrtA proteins.** For all, the spySrtA protein is shown in gray cartoon and the side chains of relevant residues are in cyan and colored by heteroatom (O=red). All distances are labeled. (**a**) Distances in the two protomers of spySrtA (PDB ID 3FN5) are not indicative of strong non-covalent interactions between the β7-β8$^{-1}$ E215 and β6$^{-2}$ T185. The lines above distances indicate that these are averages between the two rotamers of T185 in Chain A. (**b**) Because the spSrtA β7-β8 loop contains a β7-β8$^{+1}$ Glu residue, which we predict can also interact with the β6$^{-2}$ position, and the spySrtA$_{pneumoniae}$ variant is less active than the wild-type spySrtA protein (by ~20%), we *in silico* mutated the β7-β8$^{+1}$ Thr in spySrtA to Glu, T209E. We see that different rotamers of T209E reveal distances with T185 in spySrtA that are characteristic of hydrogen bonding or other non-covalent interactions.

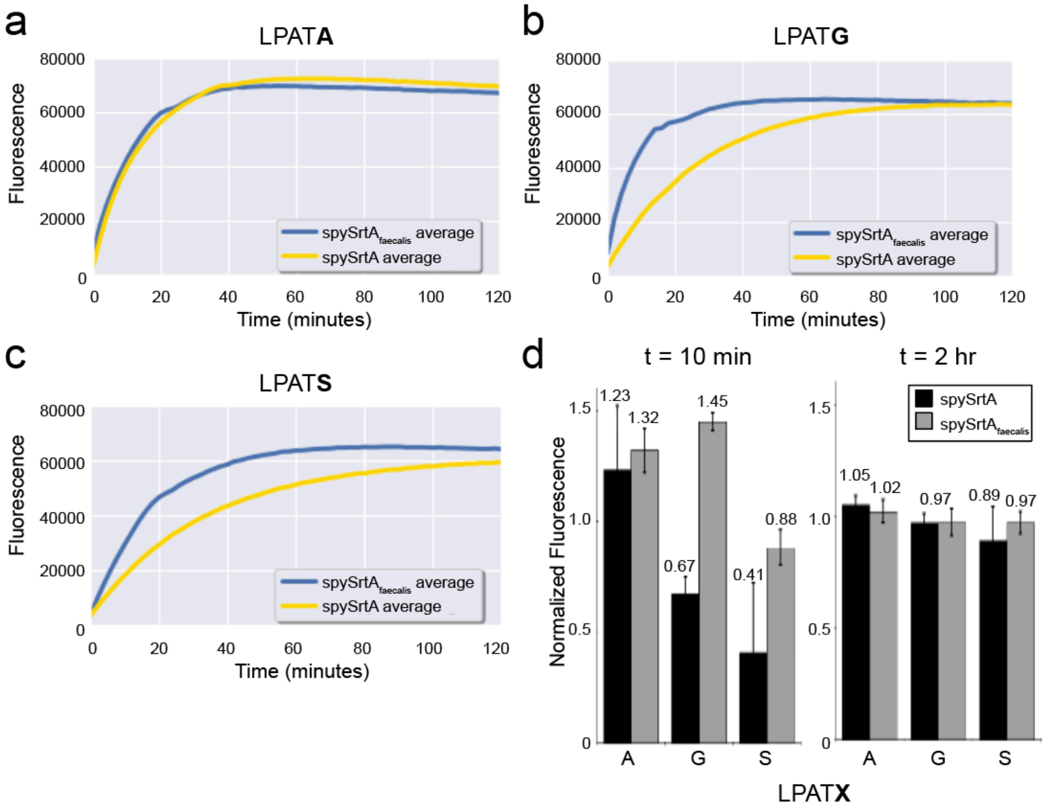


**Figure S3. Some chimeric proteins reveal differences in initial rates of the reaction.** (**a-c**) Activity traces for the fluorescence-based enzyme assays over the 2 h reaction are shown for wild-type spySrtA and spySrtA$_{faecalis}$ with the peptides: LPAT**A** (**a**), LPAT**G** (**b**), and LPAT**S** (**c**). Each plot represents the average of three or more independent experiments. (**d**) Bar graphs of average fluorescence values at t = 10 min and 2 hr for spySrtA and spySrtA$_{faecalis}$ (normalized to saSrtA:LPAT**G** control reaction at the corresponding timepoint), including standard deviation values. Average normalized values are labeled above each bar. The high error bars in the spySrtA:LPAT**A** and spySrtA:LPAT**S** reactions likely reflect the challenge in capturing early time points of these relatively fast reactions. Individual values for these reactions are: spySrtA:LPAT**A** (1.48, 0.91, and 1.30) and spySrtA:LPAT**S** (0.24, 0.24, and 0.77).

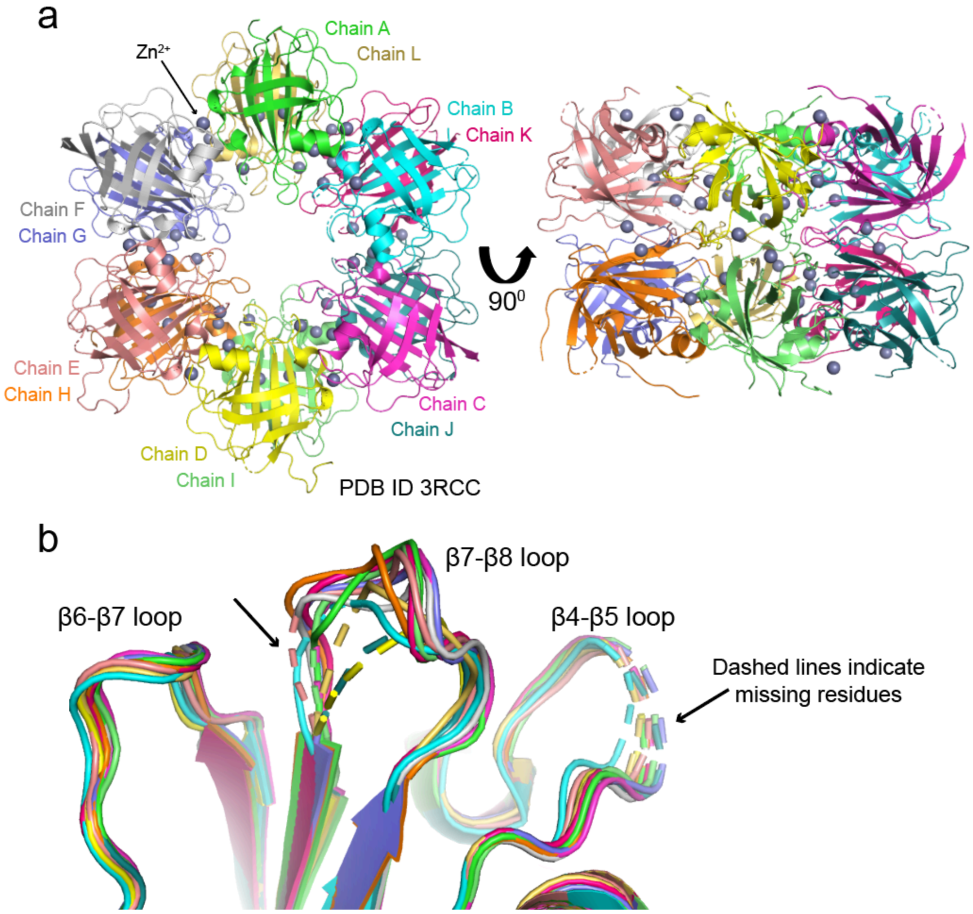


**Figure S4. The sagSrtA$_{238}$ structure.** (**a**) The dodecameric structure of the sagSrtA$_{238}$ crystal structure (PDB ID 3RCC) is shown in cartoon representation, with protomers individually colored and labeled by Chain. Zinc ions in the structure are shown as spheres. (**b**) In all protomers of the asymmetric unit in the sagSrtA$_{238}$ structure, there are missing residues in either of the β4-β5 or β7-β8 loops, or both. Colors are consistent with those in (**a**). The β4-β5, β6-β7, and β7-β8 loops are labeled and missing residues are shown as dashed lines in the cartoon.

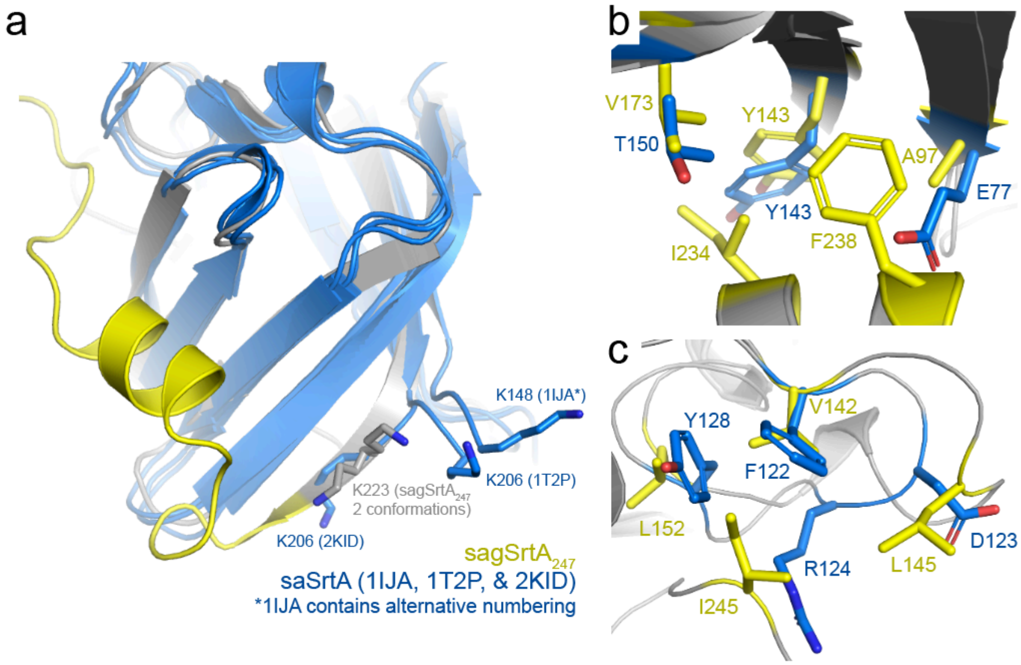


**Figure S5. Comparison of the C-termini of sagSrtA$_{247}$ and saSrtA.** In all, proteins are shown in cartoon representation, with relevant side chains as sticks and colored by heteroatom, as labeled (O=red, N=blue). (**a**) The sagSrtA$_{247}$ structure is gray, with the C-terminus colored yellow. Three saSrtA structures are aligned and colored marine blue. The residues mentioned in the main text (K223 in sagSrtA and K206, using 2KID and 1T2P numbering, in saSrtA) are shown as sticks and labeled. (**b-c**) Specific hydrophobic interactions made in sagSrtA$_{247}$ C-terminal residues with other amino acids (yellow) are not conserved in saSrtA (blue), which does not contain the additional C-terminal residues. Notably, saSrtA contains charged residues (E77, D123, R124) not present in sagSrtA. Here, only PDB ID 2KID is shown for saSrtA.

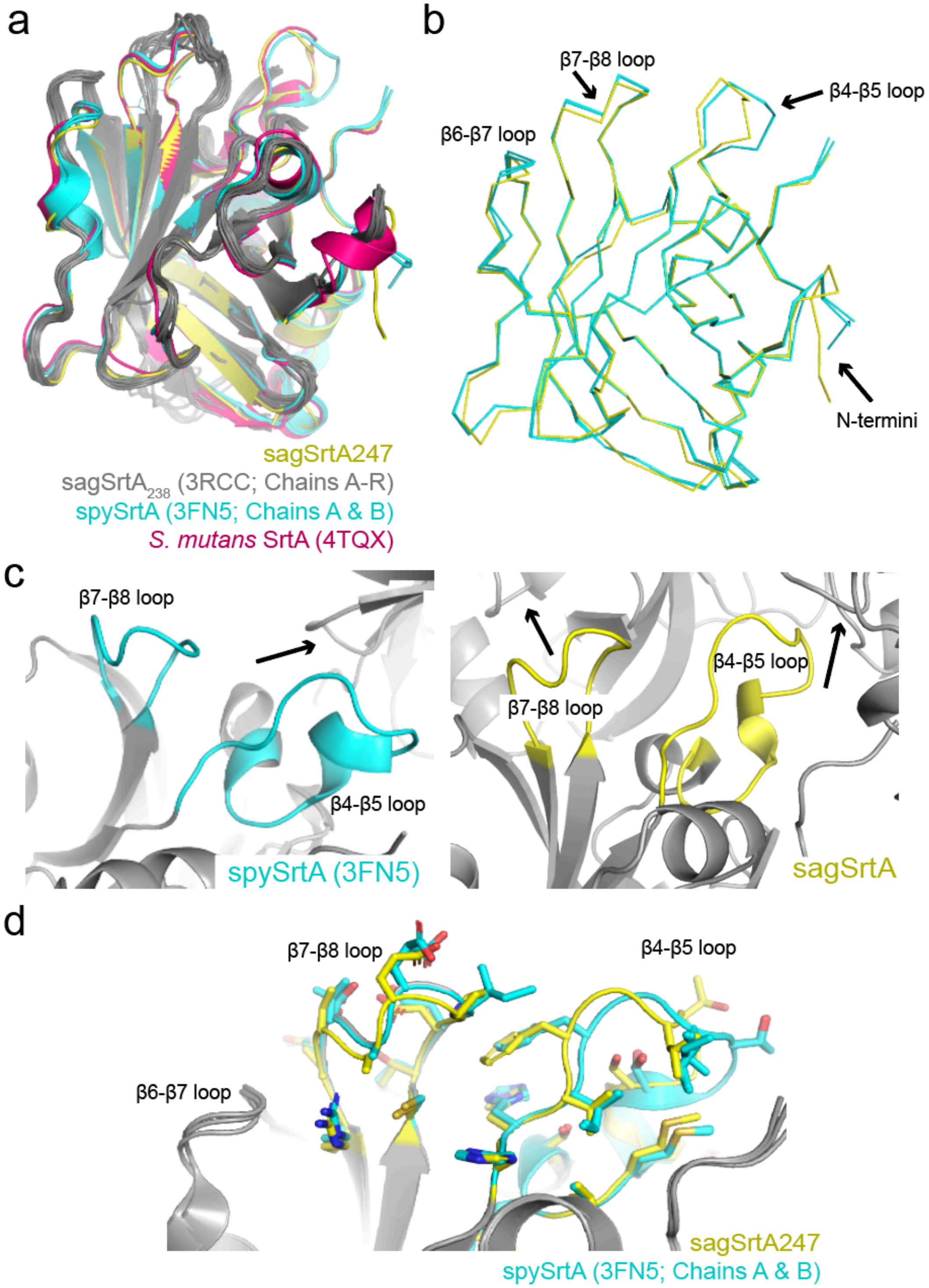


**Figure S6. The sagSrtA$_{247}$ structure is most similar to spySrtA (PDB ID 3FN5).** (**a**) Alignment of available *Streptococcus* SrtA structures with sagSrtA$_{247}$. Proteins are in cartoon representation and colored as labeled. (**b**) The sagSrtA$_{247}$ structure is most similar to the two protomers of spySrtA (3FN5). Proteins are in ribbon representation and the arrows indicate the regions of highest structural variation. (**c**) The ~1 Å displacements seen in the β4-β5 and β7-β8 loops may be due to different local crystal environments and lattice contacts. The molecules related by symmetry for spySrtA (cyan, left figure) and sagSrtA$_{247}$ (yellow, right figure) are indicated with the black arrows. (**d**) Despite the loop displacements shown in (**c**), the side chains of loop residues are in very similar locations. Side chains are shown as sticks and colored as labeled.

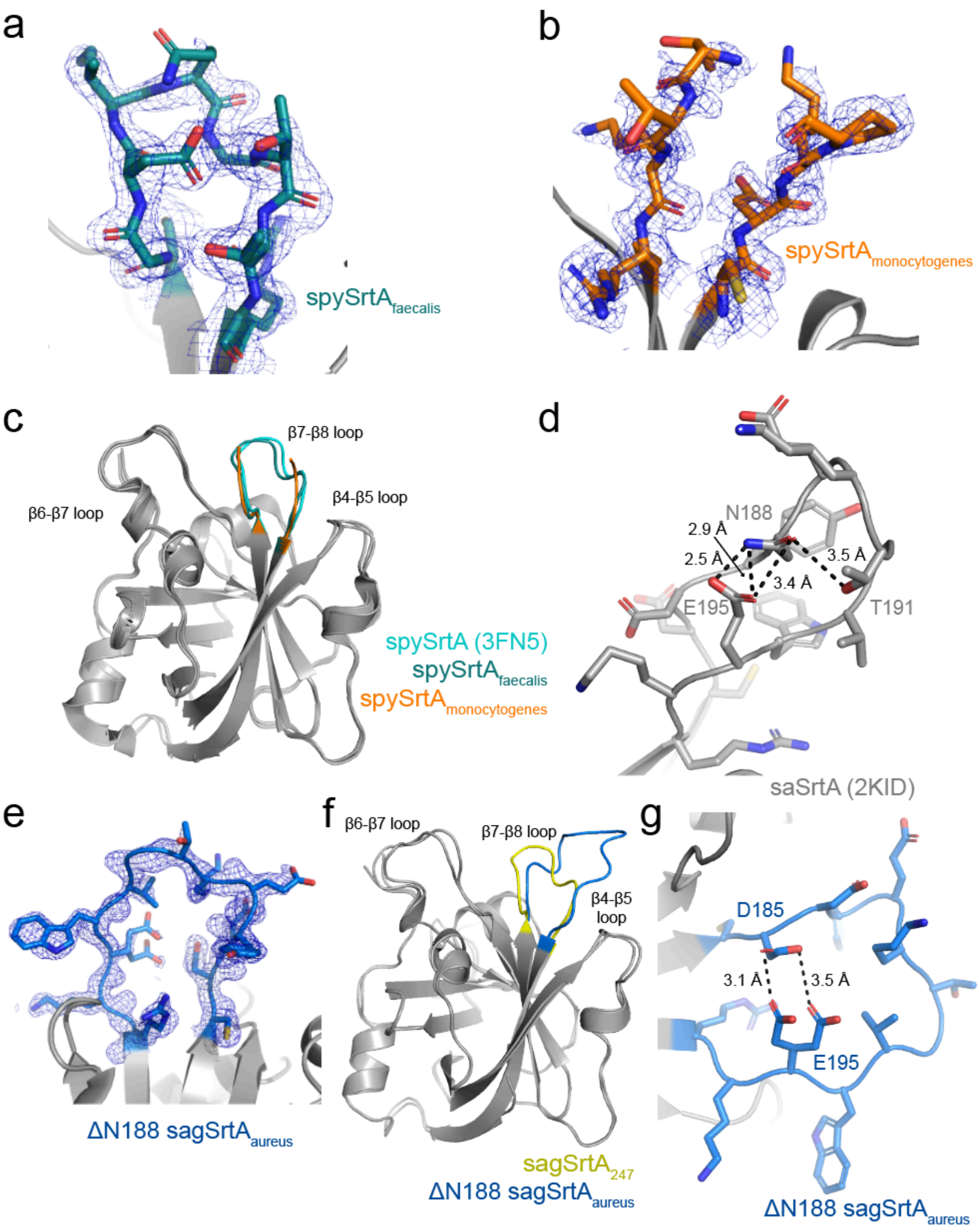


**Figure S7. Structures of sagSrtA and spySrtA β7-β8 loop chimeras.** (**a-b**) Electron density for the β7-β8 loop residues in spySrtA chimeric proteins. For all, the $2F_o$-$F_c$ electron density maps are rendered at 1σ. The structures are spySrtA$_{faecalis}$ (**a**) and spySrtA$_{monocytogenes}$ (**b**). (**c**) Alignment of the β7-β8 loop chimera structures with the respective spySrtA wild-type protein. All proteins are in cartoon representation and the β7-β8 loops are colored as labeled. (**d**) Comparison of the intraloop interactions in saSrtA (PDB ID 2KID, left figure), gray cartoon and side chains colored by heteroatom. Distances are labeled. (**e**) Electron density for ΔN188 sagSrtA$_{aureus}$, rendered as in (**a-b**). (**f**) Alignment of ΔN188 sagSrtA$_{aureus}$ with sagSrtA$_{247}$, rendered as in (**c**). (**g**) Comparison of intraloop interactions in ΔN188 sagSrtA$_{aureus}$ (right figure), marine blue cartoon and side chains colored by heteroatom, rendered as in (**d**).

**Table S2.** The β7-β8 loop sequences of *Streptococcus* proteins.

| Organism | UniProt ID | Sequence | Source |
|---|---|---|---|
| *Streptococcus_acidominimus* | A0A1Q8EFZ1_STRAI | CEDLEATMR | UniProt |
| *Streptococcus_agalactiae* | | CTDPEATER | PDB ID 3RCC; UniProt ID SRTA_STRA3 |
| *Streptococcus_alactolyticus* | A0A6N7X4S4_STRAY | CTDQAATNR | UniProt |
| *Streptococcus_alactolyticus* | A0A6N7X7P2_STRAY | CDDLQASNR | UniProt |
| *Streptococcus_anginosus* | A0A2T0FSR1_STRAP | CEDAAATSR | UniProt |
| *Streptococcus_australis* | A0A4Q1ZK73_9STRE | CEDAAATYR | UniProt |
| *Streptococcus_azizii* | A0A1V2SN59_9STRE | CEDIQATMR | UniProt |
| *Streptococcus_chenjunshii* | A0A372KPW2_9STRE | CTDAEATNR | UniProt |
| *Streptococcus_chosunense* | A0A3B0BM94_9STRE | CEDAAATER | UniProt |
| *Streptococcus_cristatus* | A0A5B0DQ60_STRCR | CEDAAATSR | UniProt |
| *Streptococcus_cuniculi* | A0A1Q8E9Q8_9STRE | CEDMDATRR | UniProt |
| *Streptococcus_danieliae* | A0A7Z0M729_9STRE | CEDAAAVYR | UniProt |
| *Streptococcus_gallolyticus* | A0A368UG76_9STRE | CTDQEATER | UniProt |
| *Streptococcus_gordonii* | A0A1V0H3H5_STRGN | CEDAAATSR | UniProt |
| *Streptococcus_gwangjuense* | A0A387AXL2_9STRE | CEDAAATER | UniProt |
| *Streptococcus_halitosis* | A0A3R8NWZ1_9STRE | CVDYNATER | UniProt |
| *Streptococcus_lutetiensis* | A0A7T9MJK1_9STRE | CTDQQATER | UniProt |
| *Streptococcus_macedonicus* | A0A081JI37_STRMC | CMDQDATER | UniProt |
| *Streptococcus_minor* | A0A3P1VGS9_9STRE | CEDAEATMR | UniProt |
| *Streptococcus_mitis* | A0A081SCL7_STRMT | CVDYDATER | UniProt |
| *Streptococcus_parasanguinis* | A0A414CL89_STRPA | CEDAAATYR | UniProt |
| *Streptococcus_parauberis* | A0A1S1ZQL8_9STRE | CTDAEATER | UniProt |
| *Streptococcus_pasteurianus* | A0A135YLJ4_9STRE | CMDQEATER | UniProt |
| *Streptococcus_periodonticum* | A0A3S9MRJ1_9STRE | CEDAAATSR | UniProt |
| *Streptococcus_pluranimalium* | A0A2L0D4A6_9STRE | CTDAEATER | UniProt |
| *Streptococcus_pneumoniae* | | CEDLAATER | Nikghalb et al. 2018. Chembiochem. PMID: 29124839 |
| *Streptococcus_pyogenes* | | CTDIEATER | PDB ID 3FN5 |
| *Streptococcus_oralis* | | CVDYNATER | Nikghalb et al. 2018. Chembiochem. PMID: 29124839 |
| *Streptococcus_ovuberis* | A0A7X6S1K7_9STRE | CEDAGAEFR | UniProt |
| *Streptococcus_rubneri* | A0A4Z1DTF3_9STRE | CDDLAATNR | UniProt |
| *Streptococcus_ruminantium* | A0A2Z5U3B1_9STRE | CTDYYATER | UniProt |
| *Streptococcus_salivarius* | A0A2W1I9H1_STRSL | CTDAEATQR | UniProt |
| *Streptococcus_suis* | | CTDYYATQR | Nikghalb et al. 2018. Chembiochem. PMID: 29124839 |
| *Streptococcus_symci* | A0A501P9G5_9STRE | CEDAAATER | UniProt |
| *Streptococcus_ursoris* | A0A7X9QHF2_9STRE | CTDAGATAR | UniProt |
| *Streptococcus_vestibularis* | A0A3E4X8P5_STRVE | CTDAEATQR | UniProt |
| *Streptococcus_xiaochunlingii* | A0A501VWP7_9STRE | CEDAAATYR | UniProt |

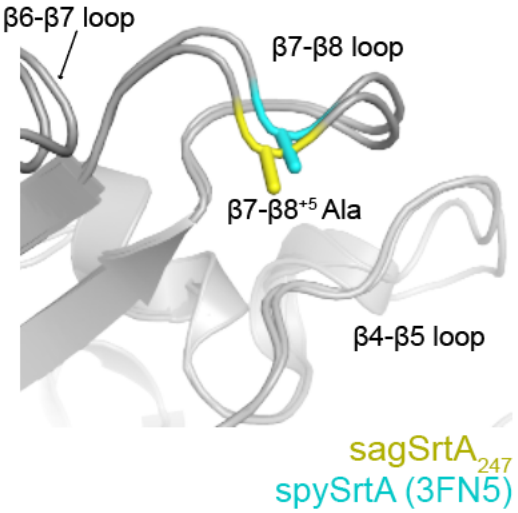


**Figure S8. A conserved alanine in the β7-β8 loops of *Streptococcus* proteins.** The location of the β7-β8$^{+5}$ Ala in the sagSrtA$_{247}$ and spySrtA structures. Both proteins are shown in gray cartoon representation with the alanine side chains shown as sticks and colored as labeled.
